# The ATPase mechanism of myosin 15, the molecular motor mutated in DFNB3 human deafness

**DOI:** 10.1101/2020.06.17.155424

**Authors:** Fangfang Jiang, Yasuharu Takagi, Arik Shams, Sarah M. Heissler, Thomas B. Friedman, James R. Sellers, Jonathan E. Bird

## Abstract

Cochlear hair cells possess an exquisite bundle of actin-based stereocilia that detect sound. Unconventional myosin 15 (MYO15A) traffics and delivers critical molecules required for stereocilia development and is essential for building the mechanosensory hair bundle. Mutations in the human *MYO15A* gene interfere with stereocilia trafficking and cause hereditary hearing loss, DFNB3. To understand the molecular mechanism of how MYO15A delivers proteins within stereocilia, we performed a kinetic study of the ATPase motor domain to characterize its mechano-chemical cycle. Using the baculovirus-Sf9 system, we purified a recombinant minimal motor domain (S1) by co-expressing the mouse MYO15 ATPase, essential and regulatory light chains that bind its IQ domains, and UNC45 and HSP90A chaperones required for correct folding of the ATPase. MYO15 purified with either UNC45A or UNC45B co-expression had similar ATPase activities (*k*_cat_ = ~ 6 s^−1^ at 20°C). Using stopped-flow and quenched-flow transient kinetic analyses, we measured the major rate constants describing the ATPase cycle, including ATP, ADP and actin binding, hydrolysis and phosphate release. Actin-attached ADP release was the slowest measured transition (~ 12 s^−1^ at 20°C), although this did not rate-limit the ATPase cycle. The kinetic analysis shows the MYO15 motor domain has a moderate duty ratio (~ 0.5) and weak thermodynamic coupling between ADP and actin binding. This is consistent with MYO15 being adapted for strain sensing as a monomer, or processive motility if oligomerized into ensembles. Our kinetic characterization enables future studies into how deafness-causing mutations affect MYO15 and ultimately disrupt stereocilia trafficking necessary for normal hearing.

## Introduction

Unconventional myosin 15, encoded by the *MYO15A* gene in humans and *Myo15* in mouse, is a member of the myosin superfamily of P-loop ATPases that generate force on actin filaments (Houdusse and Sweeney, 2016). MYO15 is expressed by hair cells of the inner ear (Liang et al., 1999; Anderson et al., 2000; Belyantseva et al., 2003) and is necessary for the structural integrity of actin-based mechanosensory stereocilia that elongate from the surface of hair cells to detect sound and accelerations (Probst et al., 1998; Belyantseva et al., 2003; Barr-Gillespie, 2015). MYO15 isoforms traffic to the distal tips of stereocilia (Belyantseva et al., 2003; Rzadzinska et al., 2004; Fang et al., 2015), which is the primary site of actin filament polymerization (Schneider et al., 2002; Zhang et al., 2012; Drummond et al., 2015; Narayanan et al., 2015). Mutations in *MYO15A* cause autosomal recessive hearing loss DFNB3 in humans (Friedman et al., 1995; Wang et al., 1998; Rehman et al., 2016), highlighting its critical function in sensory function and the need to understand how this molecular motor operates within stereocilia.

Cochlear hair cells express multiple protein isoforms of *MYO15A* / *Myo15* created through alternative mRNA splicing (Liang et al., 1999; Belyantseva et al., 2003; Fang et al., 2015). A short isoform (MYO15-2, also called MYO15-S), contains the core ATPase ‘motor domain’ and three light chain binding domains (LCBD) that serve to amplify structural changes within the motor and generate the power stroke (Fig. 1A). LCBD sites 1 and 2 are consensus IQ domains that bind to non-muscle regulatory (RLC, MYL6) and essential light chains (ELC, MYL12B), respectively (Bird et al., 2014). A third LCBD has distant similarity to the IQ consensus sequence, however no associated light chain has been reported (Bird et al., 2014). The C-terminal tail of MYO15-2 contains a Src homology 3 (SH3) domain, and similar to unconventional class VII and X myosins, myosin tail homology 4 (MyTH4) and band 4.1, ezrin, radixin, moesin (FERM) domains, in addition to a PDZ ligand at the C-terminus (Liang et al., 1999; Belyantseva et al., 2005). A larger isoform (MYO15-1, also called MYO15-L) differs from MYO15-2 solely by the addition of a 133 kDa N-terminal domain that is encoded by inclusion of a single exon in the transcript (Liang et al., 1999)(Fig. 1A). No functional domains have been identified within the proline-rich N-terminal domain, however it is essential for hearing (Fang et al., 2015).

**FIGURE 1.**
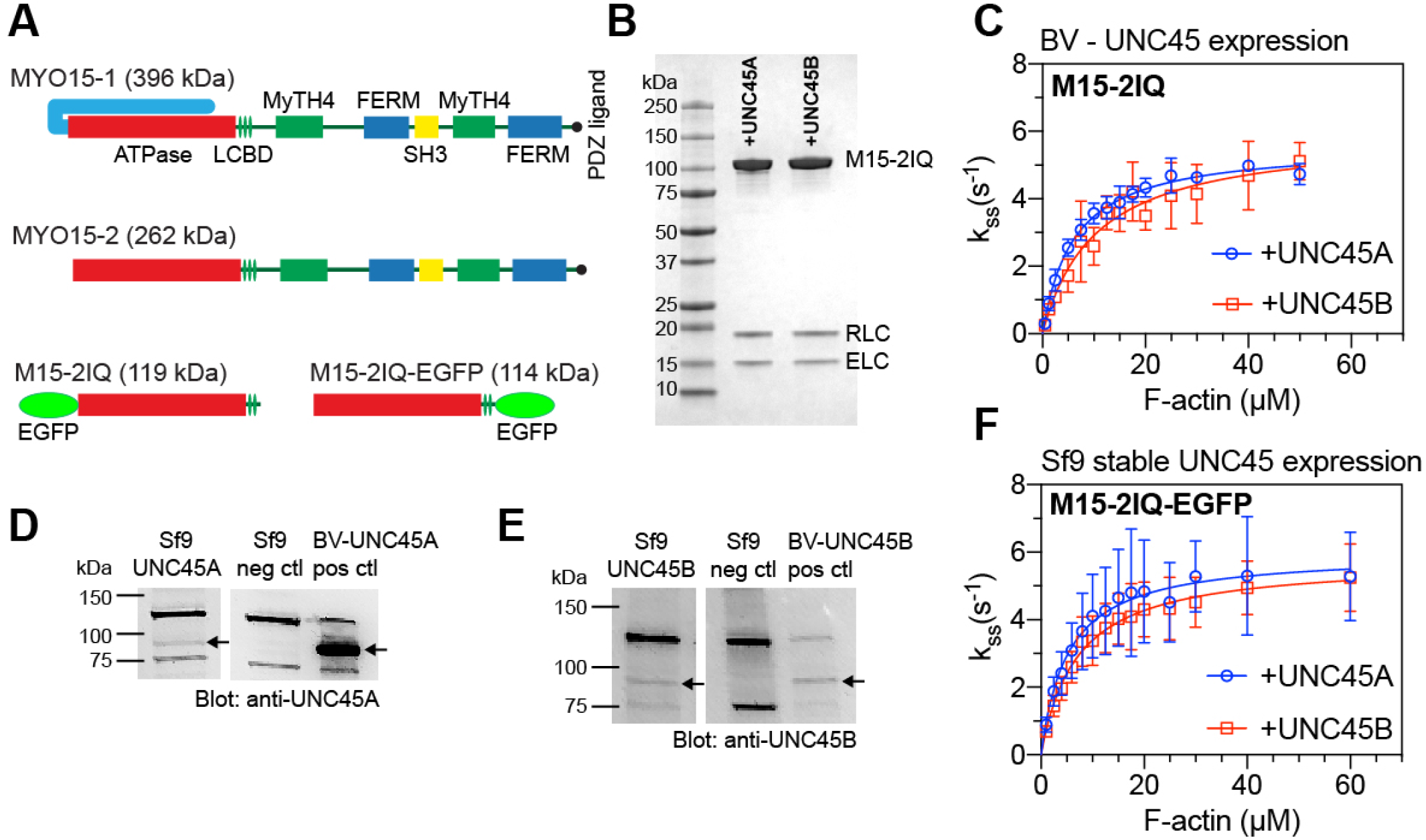
Purification and steady-state activity of the unconventional myosin 15 (MYO15) motor domain. **A)** Drawing of alternative splice isoforms of MYO15 expressed in the inner ear. Each isoform shares a common ATPase domain (red) and three light chain binding domains (LCBDs, IQ domains), in addition to MyTH4 (green), SH3 (yellow) and FERM (blue) domains. Both isoforms are identical, except for the addition of a 133 kDa N-terminal domain in MYO15-1. Enhanced green fluorescent protein (EGFP) – tagged baculovirus expression constructs used in this study are shown. **B)** SDS-PAGE analysis of the truncated motor domain (M15-2IQ) purified from Sf9 cells co-expressing baculovirus-expressed HSP90AA1, UNC45A or UNC45B, plus ELC (MYL6) and RLC (MYL12) light chains. **C)** Steady-state ATPase activation of M15-2IQ by actin filaments measured using the NADH assay. Co-expression of M15-2IQ with UNC45A (blue) yielded *k*_cat_ = 5.6 ± 0.1 s^−1^, *K*_ATPase_ = 6.1 ± 0.5 μM. Co-expression of M15-2IQ with UNC45B (red) yielded *k*_cat_ = 5.9 ± 0.4 s^−1^, *K*_ATPase_ = 10.4 ± 2.2 μM. Data are from 3 independent myosin preparations. **D,E)** Western blotting of cell lysates from *Sf9* cells engineered to stably express UNC45A (D), or UNC45B (E). Wild-type *Sf9* cells lysates are included as negative controls, and *Sf9* cells infected with either UNC45A, or UNC45B baculovirus as positive controls. Bands specific for UNC45A + UNC45B are marked with arrows. **F)** Steady-state ATPase activation of M15-2IQ-EGFP purified from *Sf*9-UNC45A, or *Sf*9-UNC45B cell lines, using the NADH assay. M15-2IQ-EGFP purified from *Sf*9-UNC45A (blue) yielded *k*_cat_ = 6.0 ± 0.5 s^−1^, *K*_ATPase_ = 5.2 ± 1.5 μM. M15-2IQ-EGFP purified from *Sf*9-UNC45B (red) yielded *k*_cat_ = 5.8 ± 0.3 s^−1^, *K*_ATPase_ = 7.0 ± 1.3 μM. Data are from 3 independent determinations, using 2 independent myosin preparations. All ATPase measurements performed at 20°C in 10 mM MOPS, 10 mM KCl, 5 mM MgCl_2_, 0.1 mM EGTA, 2 mM MgATP, 40 U•mL^−1^ LDH, 200 U•mL^−1^ PK, 1 mM PEP, 200 μM NADH. The basal ATPase activity of myosin was subtracted from each data point.

MYO15-1 and MYO15-2 have independent functions regulating stereocilia architecture in cochlear hair cells. MYO15-2 is sufficient to drive stereocilia elongation during development (Belyantseva et al., 2005; Fang et al., 2015), and in heterologous cells lines is able to traffic along filopodia similar to class VII and X myosins (Berg and Cheney, 2002; Belyantseva et al., 2003, 2005; Kerber et al., 2009; Sakai et al., 2011; Baboolal et al., 2016). MYO15-2 complexes with at least four proteins, including: whirlin (WHRN), epidermal growth factor receptor kinase substrate 8 (EPS8), guanine nucleotide-binding protein Gi subunit alpha (GNAI3) and G-protein signaling modulator 2 (GPSM2). MYO15-2 traffics this ‘elongation complex’ along filopodia actin filaments *in vitro*, and *in vivo* delivers these proteins to the tip compartment of stereocilia, where they are required for polymerization and elongation of the stereocilia actin core (Mburu et al., 2003; Belyantseva et al., 2005; Delprat et al., 2005; Manor et al., 2011; Zampini et al., 2011; Tarchini et al., 2016; Mauriac et al., 2017; Tadenev et al., 2019). The large MYO15-1 isoform is dispensable for stereocilia elongation, but is necessary for maintaining the adult length of stereocilia with active mechano-electric transduction (MET) channels (Fang et al., 2015). MYO15-1 does not traffic WHRN, EPS8, GNAI3 or GPSM2, and its associated cargo proteome is currently unknown (Fang et al., 2015). The ability of MYO15 isoforms to traffic within stereocilia is thus critical for mechano-sensory function, but their mechanisms of motility are poorly understood.

In most members of the myosin superfamily, the motor domain generates force by reversibly binding to actin filaments and undergoing an ATP-dependent mechano-chemical cycle. Kinetic tuning of the motor domain significantly diversifies motor behavior on actin filaments and allows myosins to engage in highly specific cellular functions (De La Cruz and Ostap, 2004; Nyitrai and Geeves, 2004; Bloemink and Geeves, 2011; Heissler and Sellers, 2016). Kinetic tuning of the MYO15 motor domain is currently unknown, yet this information is critical to decipher its function within stereocilia. We previously purified the motor domain of mouse MYO15 and showed it was a fast, high-duty ratio motor that moves toward the barbed end of actin filaments (Bird et al., 2014). In the present study, we have characterized the motor domain in detail using transient-state kinetic analyses to reveal its key enzymatic adaptations. Our results show that the MYO15 motor domain has characteristics consistent with sensing strain as a monomer, and that if oligomerized into an ensemble, would be sufficient to enable processive movement along actin filaments.

## Results

### UNC45A and UNC45B each promote folding of the MYO15 motor domain

UNC45 is required for the correct folding and assembly of muscle thick filaments (Lee et al., 2014), and acts as a HSP90 dependent co-chaperone that folds the muscle myosin motor domain (Barral et al., 2002; Liu et al., 2008). We and others have reported that UNC45 also catalyzes the folding of some non-muscle myosin motor domains when expressed in the *Sf9* system (Bird et al., 2014; Bookwalter et al., 2014), indicating that UNC45 acts more broadly to fold myosin motors. Mammals express two UNC45 paralogs, with 57% amino acid identity in humans. UNC45A is expressed ubiquitously throughout the body, whilst UNC45B is more restricted to striated muscle tissues although not solely expressed there (Price et al., 2002). Interestingly, cochlear hair cells express both UNC45A and UNC45B (https://umgear.org) raising the possibility that both co-chaperones might be utilized to fold MYO15 *in vivo*.

Our previous work demonstrated that co-expression of UNC45B significantly increased the yield of purified, enzymatically active MYO15 motor domain when produced in *Sf9* insect cells (Bird et al., 2014). We tested whether UNC45A could catalyze folding of the MYO15 motor domain similar to the action of UNC45B. To do this, a minimal motor domain construct (S1) of mouse MYO15 (abbreviated M15-2IQ) that contained the ATPase plus two light chain binding domains (LCBD) was expressed in *Sf9* insect cells using the baculovirus expression system. An enhanced green fluorescent protein (EGFP) moiety and FLAG epitope were fused to the N-and C-terminus of M15-2IQ, respectively (Fig. 1A). Regulatory (RLC) and essential (ELC) light chains were co-expressed to bind the LCBD (Bird et al., 2014). In addition to MYO15 and light chains, Sf9 cells were also infected with a dual-promoter baculovirus expressing HSP90AA1 and either UNC45A or UNC45B (see Experimental Procedures). Co-expression of either UNC45 paralog resulted in qualitatively similar yields of purified M15-2IQ (Fig. 1B), indicating that both catalyzed folding of the motor domain in *Sf*9 cells.

We next hypothesized that the activity of the motor domain might be influenced by the specific UNC45 chaperone recruited to fold MYO15. To test this, the steady-state ATPase activity of M15-2IQ, purified from *Sf9* cells co-expressing either UNC45A or UNC45B, was measured using an enzyme-linked NADH assay (Fig. 1C). The apparent affinity of M15-2IQ for actin in the presence of ATP is strongly dependent upon salt concentration (Bird et al., 2014), and our assays were performed with 10 mM KCl to increase the affinity of M15-2IQ for actin. ATPase rates were fit to a hyperbola to estimate *k*_cat_ = 5.6 ± 0.1 s^−1^ and *K*_ATPase_ = 6.1 ± 0.5 μM for M15-2IQ purified from *Sf*9 cells co-expressing UNC45A. The *k*_cat_ reflects the maximum catalytic activity, whilst *K*_ATPase_ is the concentration of actin required for half-maximal activation of ATPase activity. Identical measurements were performed for M15-2IQ purified from *Sf*9 cells expressing UNC45B, yielding *k*_cat_ = 5.9 ± 0.4 s^−1^ and *K*_ATPase_ = 10.4 ± 2.2 μM. ATPase parameters are summarized in Table II. We conclude that co-expression of either UNC45 paralog is sufficient to produce enzymatically active M15-2IQ, and that the use of UNC45A vs. UNC45B did not affect the overall ATPase activity. To be consistent with our previous work (Bird et al., 2014), UNC45B co-expression has been used for all M15-2IQ transient kinetics experiments described in this study.

### Development of *Sf*9-UNC45A and *Sf*9-UNC45B cell lines to streamline MYO15 motor domain purification

To simplify expression of the MYO15 motor domain, and potentially other myosin motor domains that require UNC45 to fold, we developed clonal *Sf9* cell lines that constitutively express UNC45A or UNC45B (see Experimental Procedures). Wild-type *Sf9* cells do not natively express either of these chaperones, but do express HSP90 orthologs (Carinhas et al., 2011; Bird et al., 2014). Total protein extracts from clonal *Sf*9-UNC45A and *Sf*9-UNC45B cell cultures were screened by SDS-PAGE and western blotting to confirm the expression of UNC45A or UNC45B, respectively (arrows in left lanes, Fig. 1D,E). Wild-type *Sf9* cells lysates were used as negative controls. *Sf9* cells infected with baculovirus to overexpress UNC45A / HSP90AA1, or UNC45B / HSP90AA1 served as positive controls (arrows in right lane, Fig. 1D,E). To test if these cell lines were functional for expressing MYO15, *Sf*9-UNC45A and *Sf*9-UNC45B cells were infected with baculovirus encoding the motor domain with a C-terminal EGFP and FLAG moiety (Fig. 1A, M15-2IQ-EGFP). *Sf*9 cells were additionally co-infected with baculovirus encoding the ELC and RLC light chains. We were able to purify similar quantities of M15-2IQ-EGFP from both *Sf*9-UNC45A and *Sf*9-UNC45B cells. Purified M15-2IQ-EGFP from either cell line had similar actin-activated ATPase activities. M15-2IQ-EGFP purified from *Sf*9-UNC45A cells exhibited *k*_cat_ = 6.0 ± 0.5 s^−1^ and *K*_ATPase_ = 5.2 ± 1.5 μM, whilst M15-2IQ-EGFP purified from Sf9-UNC45B had *k*_cat_ = 5.8 ± 0.3 s^−1^ and *K*_ATPase_ = 7.0 ± 1.3 μM (see Table II). These results corroborate our findings that the ATPase activity of MYO15 does not differ between usage of UNC45A vs. UNC45B, and further shows that the *Sf*9-UNC45A and *Sf*9-UNC45B cells lines are effective at expressing enzymatically active myosin motor domains.

### Interaction of MYO15 with ATP

To measure the kinetic and equilibrium constants describing the ATPase cycle of M15-2IQ (see Scheme I), we performed a series of pre-steady state experiments using stopped-flow and quenched flow techniques. ATP generated a robust increase in intrinsic M15-2IQ protein fluorescence excited at 297 nm in a stopped-flow spectrophotometer, and we used this property to measure the ATP binding mechanism. M15-2IQ (0.25 μM) was rapidly mixed under pseudo-first order conditions with increasing concentrations of [ATP] ranging up to 5 mM. Experimentally recorded fluorescence transients were fit to a mono-phasic exponential increase (Fig. 2A), and observed rate constants (*k*_obs_) varied hyperbolically with respect to [ATP] (Fig. 2B). This reaction was modeled using a two-step binding mechanism (Scheme 2), where ATP and myosin form a collision complex in rapid equilibrium (*K*_1_), followed by an isomerization (*k*_+2_) to an enhanced fluorescence state (M*•ATP).

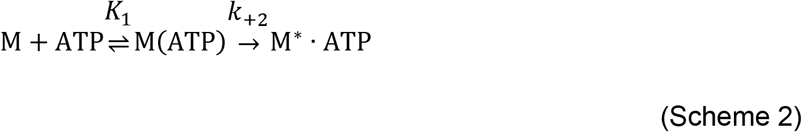

**FIGURE 2.**
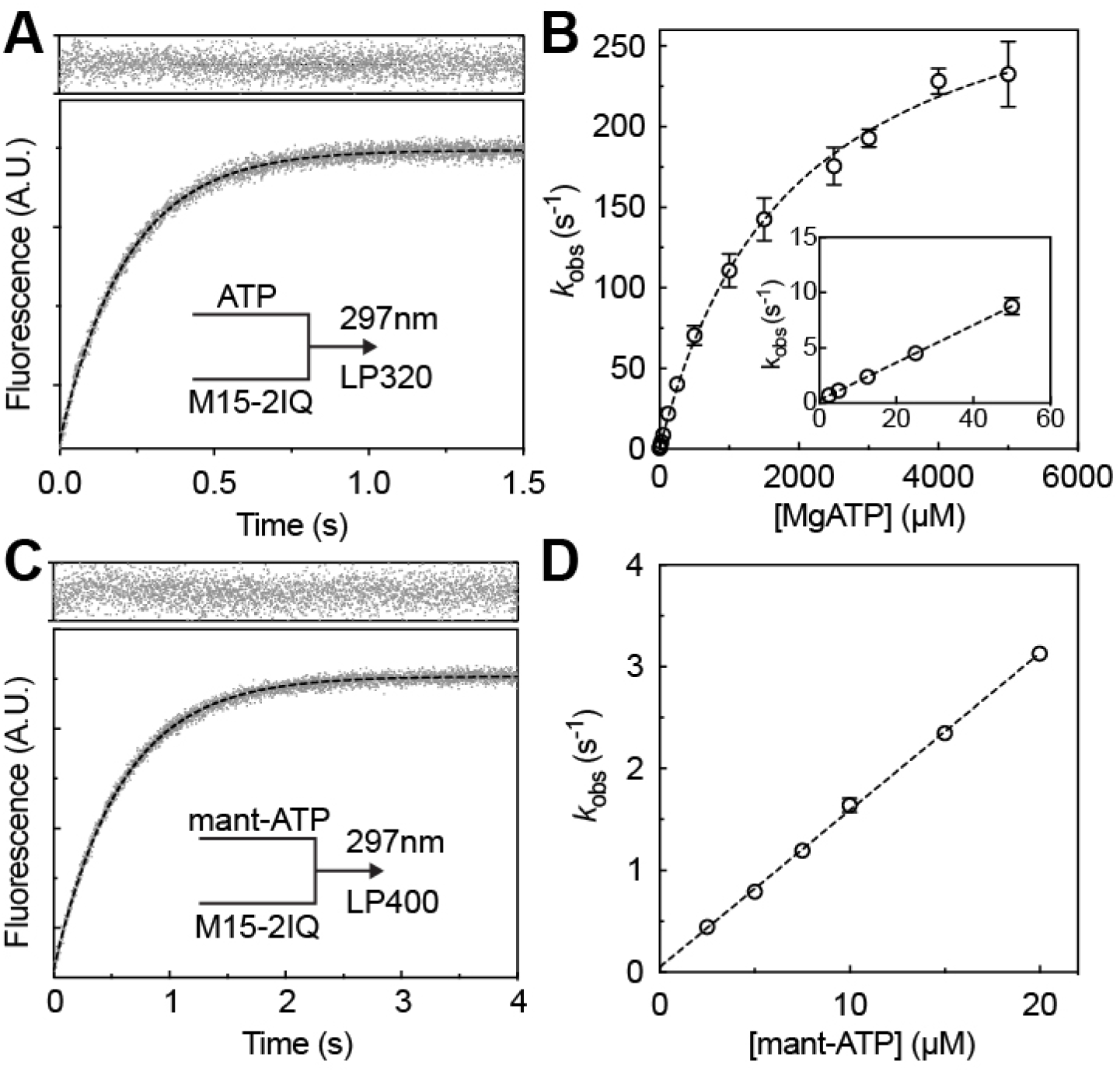
Transient kinetic analysis of ATP binding to M15-2IQ using stopped flow fluorescence spectroscopy. **A)** Intrinsic fluorescence enhancement as 0.25 μM M15-2IQ is rapidly mixed with 25 μM ATP. The transient was fit to a single exponential equation, *I(t) = −9.2·e^−4.52t^ + C* (dotted line, residuals above). **B)** Dependence of intrinsic fluorescence observed rate constants (*k*_obs_) upon ATP concentration. The hyperbola *k*_obs_ = (*K*_1_·*k*_+2_·[ATP]) / (1 + *K*_1_·[ATP]) is fitted (dotted line), where 1/*K*_1_ = 1898 ± 145 μM, *k*_+2_ = 322 ± 10 s^−1^. Inset: *k*_obs_ is shown at lower [ATP]. ATP binding was essentially irreversible. **C)** Fluorescence enhancement of 10 μM mant-ATP after rapid mixing with 0.25 μM M15-2IQ. The mant fluorophore was excited by FRET from vicinal tryptophan residues. The transient was fit to a single exponential equation *I(t) = −5.9 e^−1.71t^ + C* (dotted line, residuals above). **D)** Dependence of *k*_obs_ upon mant-ATP concentration reveals the apparent association rate constant *K*_1_·*k*_+2_ = 0.15 ± 0.002 μM^−1^·s^−1^ with an intercept of ~ 0, showing irreversible binding of mant-ATP to M15-2IQ. For all experiments, conditions in the observation cell were 20 mM MOPS (pH 7.0), 100 mM KCl, 5 mM MgCl_2_, 0.1 mM EGTA at 20 °C. Concentrations are post-mixing in the observation cell. Experimental data were measured from 3 independent myosin preparations.

The apparent association rate constant (*K*_1_•*k*_+2_) for ATP binding was 0.17 ± 0.005 μM ^−1^s^−1^, measured from the initial gradient at low [ATP] (Fig. 2B, inset). A hyperbolic fit to the intrinsic fluorescence data yielded estimates for the ATP dissociation constant 1/*K*_1_ = 1898 ± 145 μM and isomerization rate constant *k*_+2_ = 322 ± 10 s^−1^ (Fig. 2B). Our data indicated that ATP binding was effectively irreversible within the range of measurement uncertainty (e.g. *k*_-2_ ~ 0). Mouse MYO15 has a conserved tryptophan (W432) in the relay loop, homologous to W501 in conventional myosin (MhcA, *D. discoideum*) and W512 in smooth muscle myosin (*G.gallus*). In conventional myosin, this residue is proposed to sense conformational changes occurring with ATP hydrolysis (Cremo and Geeves, 1998; Málnási-Csizmadia et al., 2000). Since the exact contribution of W432 to the intrinsic fluorescence signal of MYO15 is unclear, we attribute the maximum observed rate to represent nucleotide binding (*k*_+2_) and the actin-detached hydrolysis rate (*k*_+3_ + *k*_-3_).

ATP binding was independently assessed using the fluorescent nucleotide analog, 2’-deoxy-mantATP (mantATP). Fluorescence resonance energy transfer (FRET) from vicinal tryptophan residues was used to excite mantATP as it bound to M15-2IQ. M15-2IQ (0.25 μM) was rapidly mixed under pseudo-first order conditions with mantATP titrated from 2.5 μM to 20 μM in the stopped flow. Fluorescence transients were well fit by a mono-phasic exponential increase (Fig. 2C), with observed rate constants (*k*_obs_) varying linearly with [ATP] (Fig. 2D). Linear regression to the observed rate constants yielded the association rate constant, *K*_1_•*k*_+2_ = 0.15 ± 0.002 μM ^−1^s^−1^, similar to that measured using intrinsic fluorescence.

We next measured ATP binding to actomyosin. ATP-induced dissociation of the acto-M15-2IQ complex was followed using the decrease of orthogonally-scattered light in a stopped-flow experiment. Nucleotide-free M15-2IQ (0.125 μM) pre-equilibrated with actin filaments (0.25 μM) was rapidly mixed with [ATP] under pseudo-first order conditions. Light scattering was followed at 340 nm to monitor dissolution of the actomyosin complex. At [ATP] < 5 μM, observed transients followed a mono-phasic exponential time-course, however at [ATP] > 12.5 μM, transients were fit to a bi-phasic exponential decay with clearly defined faster and slower rates (Fig. 3A). A two-step serial binding mechanism cannot be used to model this response, since both fast and slow observed rate constants phases saturated at higher [ATP] (Fig. 3B). Similar kinetics of ATP binding to actomyosin have been reported for MYO1B and MYO19 (Geeves et al., 2000; Lewis et al., 2006; Ušaj and Henn, 2017). Following those studies, we interpret these data according to Scheme 3, where nucleotide-free acto-M15-2IQ is in equilibrium (*K*_α_) between a nucleotide-insensitive (AM’) and sensitive (AM) state competent to bind ATP (Geeves et al., 2000). In the nucleotide sensitive state, ATP and acto-M15-2IQ form a collision complex in rapid equilibrium (*K*_1_’) that isomerizes (*k’*_+2_ + *k’*_-2_) to an AM•ATP state that rapidly dissociates (*K*_8_) from actin. Population of the A + M•ATP state results in a decrease in light scattering.

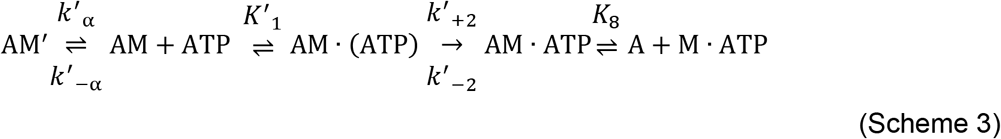

**FIGURE 3.**
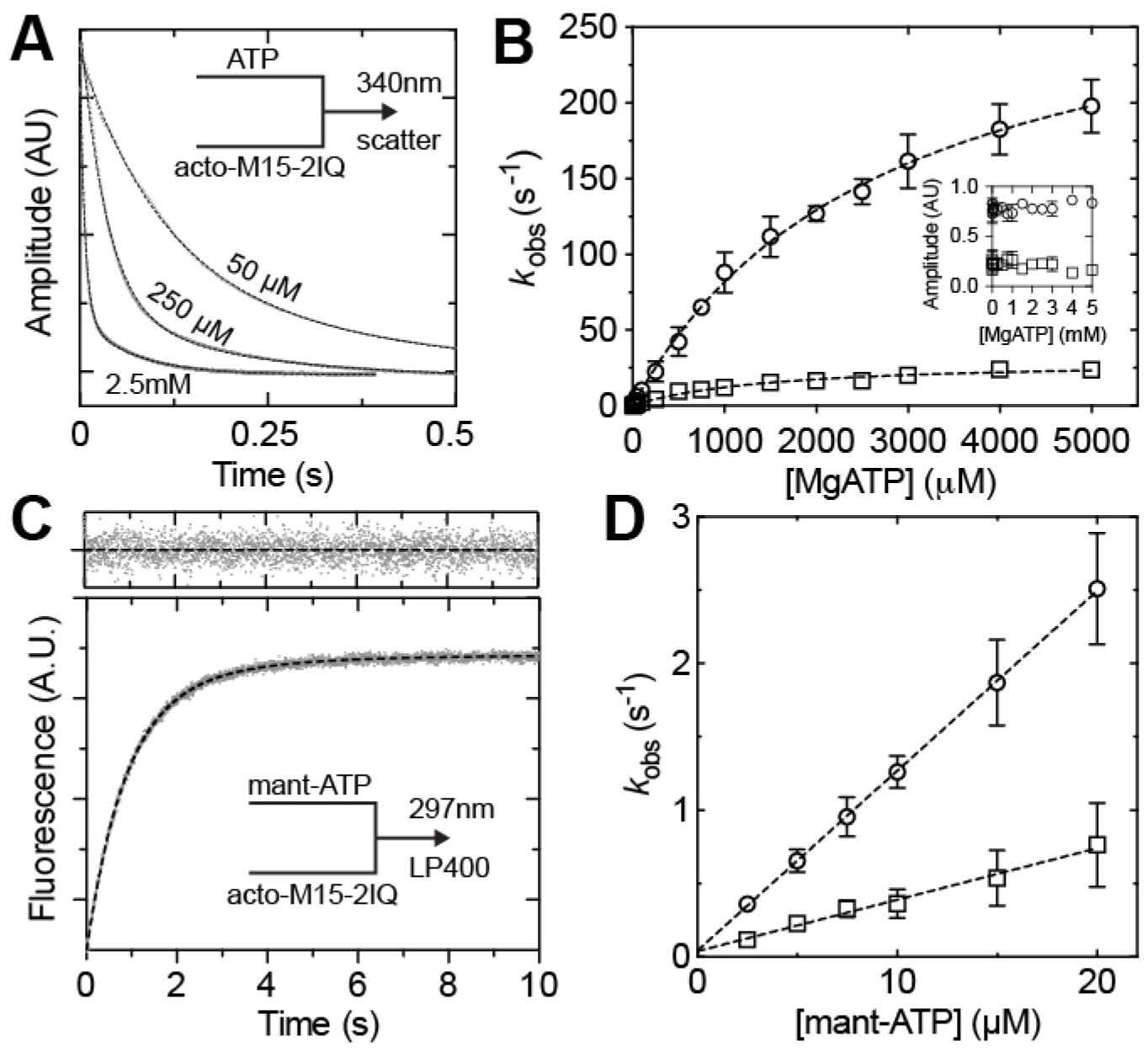
Transient kinetic analysis of ATP binding to acto-M15-2IQ using stopped flow spectroscopy. **A)** Reduction in orthogonal light scattering (measured at 340nm) as 50, 250 or 2500 μM MgATP binds to a pre-incubated mix of 0.25 μM M15-2IQ plus 0.5 μM actin in the stopped flow. Transients were fit to a bi-phasic exponential (dotted lines) with *I(t) = 36.2·e^−8.71^ + 13.3·e^−1.7t^ + C, I(t) = 42.2oe^−287^ + 10.1·e^−53^ + C*, *I(t) = 37.5oe^−1506^ + 8.4oe^−147^ + C*, for 50, 250 and 2500 μM MgATP, respectively. **B)** Dependence of fast and slow observed rate constants upon [MgATP]. Hyperbolae were fit (dotted lines) to both fast and slow phases to yield maximum values of 307 ± 15 s^−1^ and 30.9 ± 1.9 s^−1^, with half maximal activation at 2749 ± 268 μM and 1428 ± 225 μM, respectively. Inset: normalized amplitudes of fast and slow phases at different [ATP]. **C)** Fluorescence enhancement of 10 μM mant-ATP after rapid mixing with 0.25 μM M15-2IQ + 0.5 μM actin. The mant fluorophore was excited by FRET from vicinal tryptophan residues in M15-2IQ. The transient was fit to a bi-phasic exponential *I(t) = - 5.0·e^−12^ – 0.9·e^−0.4t^ + C* (dotted line, fit residuals above). **D)** Dependence of fast and slow observed rate constants upon [mant-ATP]. A linear fit to the fast phase yields the apparent association rate constant *K’*_1_·*k’*_+2_ = 0.12 ± 0.01 μM^−1^·s^−1^ with an intercept of ~ 0, showing irreversible binding of mant-ATP to acto-M15-2IQ. Conditions in the observation cell were 20 mM MOPS (pH 7.0), 100 mM KCl, 5 mM MgCl_2_, 0.1 mM EGTA at 20 °C. All concentrations are post-mixing in the observation cell. Experimental data were measured from 3 independent myosin preparations.

The fast phase (*k*_fast_) was interpreted as direct binding of ATP to the nucleotide-sensitive (AM) state (Fig. 3B), and the apparent association rate constant was estimated from the initial gradient of this response at low [ATP], *K*’_1_•*k*’_+2_ = 0.12 ± 0.02 μM ^−1^s^−1^. A hyperbolic fit to *k*_fast_ yielded the maximum isomerization rate *k’*_+2_ = 306.8 ± 15.1 s^−1^ (*k’*-2 was ~ 0 within the measurement uncertainty) and 1/*K’*_1_ = 2749 ± 268 μM ^−1^ (Fig. 3B). The slow observed phase represented *k’*_α_, the transition rate from the nucleotide-insensitive (AM’) to sensitive binding state (AM). This interpretation is valid at high [ATP], where the condition *K*’_1_•*k*’_+2_[ATP] >> *k’*_-α_ is satisfied. We estimated the maximum rate *k’α* = 30.9 ± 1.9 s^−1^ using a hyperbolic fit to the *k*_slow_ second-order response (Fig. 3B). The equilibrium constant (*k’*_α_) was estimated from the relative amplitudes of the fast and slow phases, which represent the respective fraction of acto-M15-2IQ molecules in nucleotide sensitive / insensitive states, *K’*_α_ = A_fast_ / A_slow_ = 3.8 ± 1.0 (Fig. 3B, inset).

ATP binding to actomyosin was independently probed using mant-ATP fluorescence excited by FRET from vicinal tryptophan residues in M15-2IQ. Nucleotide-free M15-2IQ (0.25 μM) was pre-incubated with actin filaments (0.5 μM) and rapidly mixed with mantATP under pseudo-first order conditions in the stopped flow. Similar to our light scattering measurements, fluorescent transients were well fit to a bi-phasic exponential increase (Fig. 3C) up to the maximum mant-ATP concentration probed (20 μM). Both observed rate constants varied linearly with [ATP], and a linear regression fit to the fast phase was interpreted as the apparent association rate constant of binding to the nucleotide sensitive fraction, *K’*_1_ • *k*’_+2_ = 0.12 ± 0.01 μM^−1^·s^−1^, in exact agreement with our estimate using light scattering (Fig. 3D).

### ATP hydrolysis and actin-activated phosphate release from MYO15

The rate of ATP hydrolysis by M15-2IQ in the absence of actin filaments was measured directly in a multiple-turnover quenched flow experiment. M15-2IQ (3 μM) was rapidly mixed under pseudo-first order conditions with 0.5 mM [gamma-^32^P]-ATP and aged for increasing periods of time, prior to acid-quench and determination of hydrolyzed Pi liberated during the reaction. The measured Pi-burst was well fit by a single exponential with *k*_obs_ = 50.2 ± 5.7 s^−1^ (Fig. 4A), representing the net flux of actin-detached forward and reverse hydrolysis (*k*_obs_ = *k*_+3_ + *k*_-3_). The amplitude of the Pi-burst was 0.57 mol P_i_ : mol myosin (Fig. 4A), and the equilibrium constant (*K*_3_) was calculated using the relation 0.57 = *K*_3_ / (1 + *K*_3_), *K*_3_ = 1.34 ± 0.07 (De La Cruz and Ostap, 2009). Our data indicates that M15-2IQ undergoes significant reverse hydrolysis and that the M•ATP and M•ADP•Pi states are both significantly populated at steady-state. The observed rate of hydrolysis in this multiple-turnover experiment was comparable to the rate of ATP binding measured using intrinsic fluorescence at an equivalent [ATP] (Fig. 2A,B), demonstrating that hydrolysis was likely rate-limited by ATP binding in this experiment. Although we likely did not probe the maximum rate of hydrolysis, our data shows that this transition does not rate-limit the catalytic cycle of M15-2IQ.

**FIGURE 4.**
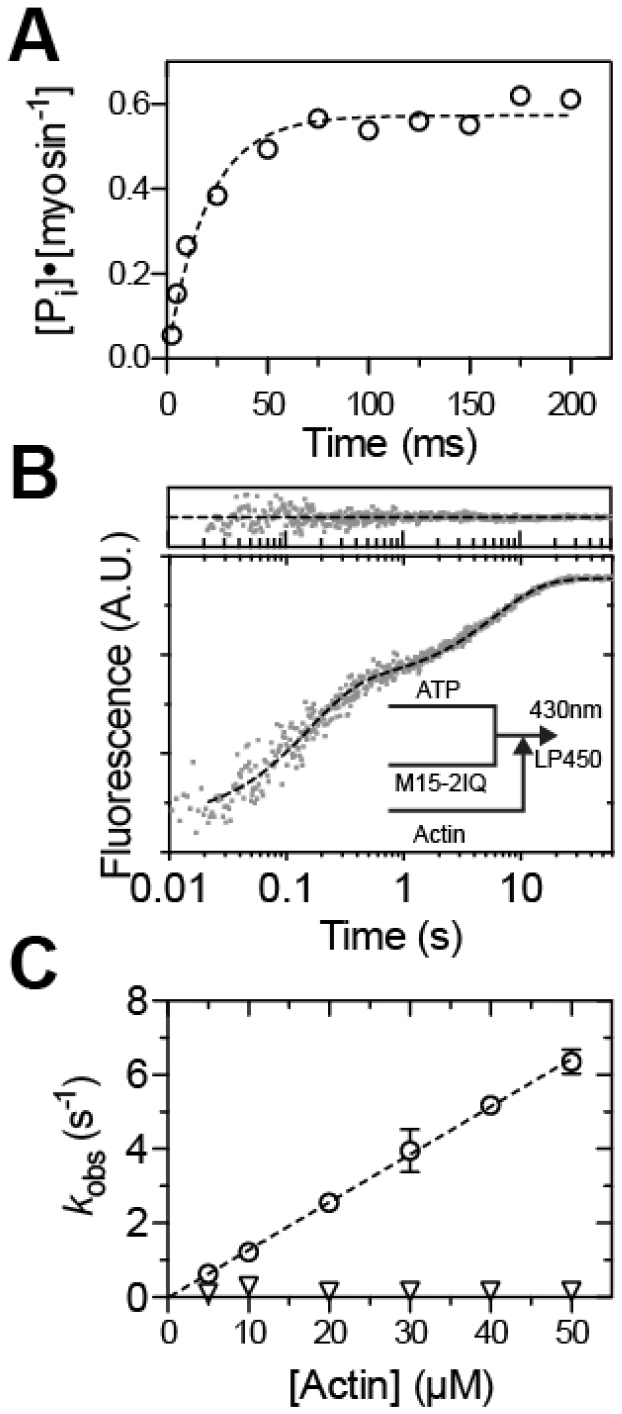
Hydrolysis and actin-activated phosphate release. **A)** ATP hydrolysis measured by quenched-flow. M15-2IQ (3 μM) was reacted under multiple turnover conditions with 0.5 mM [γ-^32^P]ATP and aged for varying intervals before acid quench. The observed phosphate burst was fit to a single exponential, *I(t) = (K_3_/1 + K_3_)·(1-e^−kobs·t^*) to yield *k*_obs_ = 50.2 ± 5.7 s^−1^ and *K*_3_ = 1.34 ± 0.07. Data are from one myosin preparation. **B)** Double-mixing stopped flow experiment to measure actin-activated phosphate release during a single turnover. M15-2IQ (2 μM) was 1:1 mixed with 1 μM ATP and aged for 5 seconds (premix concentrations), followed by a subsequent 1:1 mix with 50 μM actin (final condition in reaction cell, 1 μM myosin, 0.5 μM ATP, 50 μM actin). MDCC-PbP was included in all solutions at 5 μM. Transient is shown overlaid with *I(t) = −1.39e^−6.59t^ − 1.06e^−0.17t^ + C* (dotted line, fit residuals above). **C)** Dependence of observed rate constants upon [actin]. A linear fit to the fast phase (circles) yields *K*_9_·*k’*_+4_ = 0.13 ± 0.004 μM^−1^ s^−1^. The slow observed phase (triangles) did not vary systematically with [actin]. Data are from two independent determinations, from one myosin preparation. Conditions in the reaction loop / observation cell were 20 mM MOPS (pH 7.0), 100 mM KCl, 5 mM MgCl_2_, 0.1 mM EGTA at 20°C.

Actin-activated Pi release from M15-2IQ was examined using a single turnover, dual-mixing stopped flow experiment. M15-2IQ (2 μM) was rapidly mixed with a sub-stoichiometric quantity of ATP (1 μM) and aged for 5 seconds to allow for hydrolysis. The aged reaction was then rapidly mixed with actin filaments to accelerate Pi release from the AM•ADP•P_i_ quaternary complex. A fluorescently labeled phosphate binding protein (MDCC-PbP) was included in all solutions to monitor P_i_ release in real-time (Brune et al., 1994). In the absence of actin filaments in the second mix, P_i_ release followed a single exponential time course with *k*_obs_ = 0.08 ± 0.001 s^−1^. This rate is equivalent to the steady-state ATPase activity of M15-2IQ measured in the absence of actin (see Table II) and confirms that P_i_ release rate-limits the basal ATPase cycle in the absence of actin filaments. When actin filaments were introduced into the second mix to accelerate P_i_ release, we observed a bi-phasic exponential increase in MDCC-PbC signal (Fig. 4B). The fast-observed rate (*k*_fast_) varied linearly with [actin], whilst the slow-observed rate (*k*_slow_) was insensitive to [actin] and remained constant at ~ 0.16 s^−1^ (Fig. 4C). The fast phase was interpreted as P_i_ release upon direct binding of M•ADP•P_i_ to actin, whilst the slower phase may originate from the equilibrium of pre-hydrolysis M•ATP species that undergo actin-attached hydrolysis (M•ATP => AM•ATP => AM•ADP•P_i_), that subsequently rate-limits P_i_ release (White et al., 1997; Kovács et al., 2005; Haithcock et al., 2011). A linear regression fit to the *k*_fast_ second-order response yielded an apparent association rate constant, *K*_9_•*k’*_+4_ = 0.13 ± 0.004 μM^−1^s^−1^, indicating weak affinity of the M•ADP•P_i_ species for actin. There was no evidence of saturation in the fast-observed phase that would allow the maximum rate of P_i_ release (*k’*_+4_) to be determined. Given the fastest P_i_ release measured here (~ 6 s^−1^ at 50 μM actin) was already 4 times faster than the equivalent steady-state activity (~ 1.5 s^−1^) under identical buffer conditions at this actin concentration (Bird et al., 2014), we conclude that P_i_ release does not rate-limit the catalytic cycle of M15-2IQ in the presence of actin.

### The interaction of MYO15 with actin filaments

A stopped-flow fluorescence assay was used to measure binding of M15-2IQ to actin in the absence of ATP. We used the quenching of pyrene-iodoacetamide coupled to Cys374 of actin as a probe for the formation of the strongly-bound actomyosin state (Taylor, 1991). Nucleotide-free conditions were ensured by the pretreatment of actin and myosin solutions with apyrase. M15-2IQ (ranging from 0.05 – 0.3 μM) was rapidly mixed with an excess of pyrene actin (ranging from 0.5 – 3 μM) under pseudo first-order conditions in the stopped flow spectrophotometer. We held the myosin-to-actin ratio constant throughout the titration to ensure sufficient signal-to-noise at higher actin concentrations. The time course of pyrene fluorescence quenching was fit to a mono-phasic exponential decay (Fig. 5A) with observed rate constants (*k*_obs_) that varied linearly with respect to [actin] (Fig. 5B). These data were interpreted using a one-step binding mechanism, and linear regression fit to *k*_obs_ = *k*_+6_[actin] + *k*_-6_ yielded the apparent association rate constant *k*+_6_ = 3.18 ± 0.15 μM^−1^ s^−1^. The apparent dissociation rate *k*_-6_ = 0.65 ± 0.3 s^−1^ was measured from the y-axis intercept (Fig. 5B). These experiments were repeated in the presence of 0.5 mM ADP to saturate the M15-2IQ nucleotide binding site (apyrase was omitted in these experiments). Fluorescence transients in the presence of ADP were well modelled by a mono-phasic exponential decay, and the observed rate constant again varied linearly with [actin] (Fig. 5B). Linear regression to this response yielded the apparent association rate constant *k*_+10_ = 1.75 ± 0.09 μM^−1^ s^−1^ and apparent dissociation rate *k*-10 = 0.55 ± 0.17 s^−1^. Dissociation equilibrium constants for M15-2IQ binding to actin were calculated using the relation at equilibrium 1 / *K*_6_ = *k*_-6_ / *k*_+6_, yielding *K*_A_ = 1 / *K*_6_ = 200 ± 100 nM under nucleotide-free conditions and *K*_DA_ = 1 / *K*_1_0 = 310 ± 97 nM in the presence of saturating ADP. We conclude that M15-2IQ has a moderate affinity for actin in the absence of ATP. The ratio of actin affinity in the presence, and absence of ADP, *K*_DA_ / *K*_A_ = 1.6, indicating weak thermodynamic coupling between actin and ADP binding to M15-2IQ.

**FIGURE 5.**
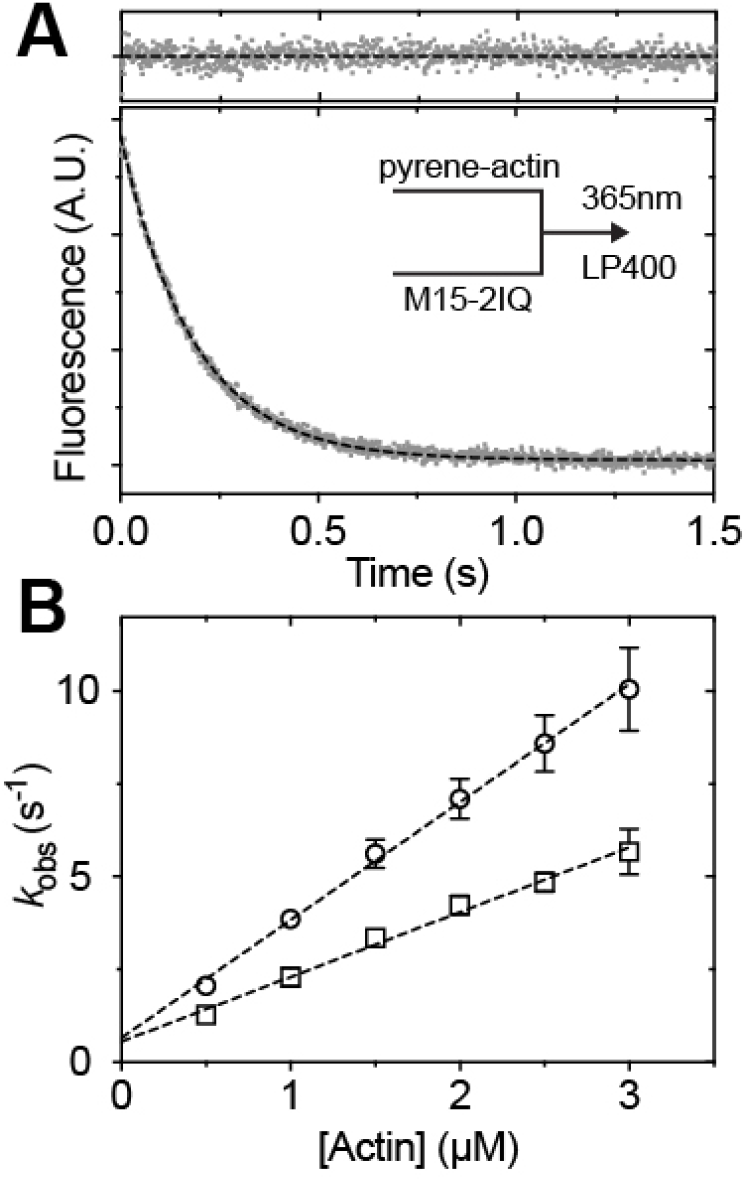
The interaction of M15-2IQ with actin filaments. **A)** Reduction in fluorescence measured as 0.15 μM M15-2IQ binds and quenches 1.5 μM pyrene-labeled actin in a stopped-flow spectrophotometer in the absence of nucleotide (apyrase-treated). The transient was fit to a single exponential function *I(t) = 5.6e^−5.4t^ + C* (dotted line, fit residuals above). Similar experiments were repeated in the presence of 1mM ADP (not shown). **B)** Dependence of *k*_obs_ upon [actin] is shown for nucleotide-free conditions (circles) and with 1 mM ADP present (squares). Linear regression to *k*_obs_ = *k*+A[actin] + *k*-A yields the association rate constant for actin binding *k*_+6_ = 3.18 ± 0.15 μM^−1^s^−1^ and off-rate *k*_-6_ = 0.65 ± 0.3 s^−1^. In the presence of ADP, *k*_+10_ = 1.75 ± 0.09 μM^−1^s^−1^ and *k*-10 = 0.55 ± 0.17 s^−1^. Experimental conditions in observation cell: 0.05 – 0.3 μM M15-2IQ, 0.5 – 3 μM pyrene-actin, 1 mM ADP (optional), 20 mM MOPS (pH 7.0), 100 mM KCl, 5 mM MgCl_2_, 0.1 mM EGTA at 20 °C. Experimental data were measured from 3 independent myosin preparations.

### ADP release from acto-MYO15 is the slowest step in the ATPase cycle

We used enhancement of intrinsic protein fluorescence to measure the kinetics of ADP binding to M15-2IQ. Although intrinsic protein fluorescence is well established to be sensitive to structural changes induced by ATP binding and hydrolysis, in muscle myosin it is also responsive to ADP binding (Morita, 1967; Werber et al., 1972). We similarly found that mixing of ADP with M15-2IQ induced a robust intrinsic fluorescence transient. M15-2IQ (0.25 μM) was rapidly mixed in the stopped flow with ADP under pseudo-first order conditions whilst monitoring intrinsic fluorescence. ADP in the reaction was titrated across a concentration range of 2.5 μM to 500 μM. At low [ADP], the observed transients followed a mono-phasic exponential increase, however at higher [ADP] an initial lag phase became apparent (Fig. 6A). We attempted to fit this transient using a double exponential function, but could not resolve the fast phase (*k*_obs1_) within acceptable error. Instead, we fit the observed transients to a single exponential (*k*_obs2_) at lower [ATP], and excluded the short lag phase from the fitting procedure at higher [ATP]. Observed rate constants for the slower phase (*k*_obs2_) varied hyperbolically with respect to [ADP] (Fig. 6B), consistent with ADP binding being at least a 2-step mechanism. We modeled the reaction as shown in Scheme 4, where myosin and ADP initially form a weakly bound state M(ADP), that isomerizes to a strongly bound M*•ADP state with enhanced fluorescence. These states are equivalent to the myosin nucleotide binding pocket in ADP-bound ‘open’, and ‘closed’ conformations, respectively (Nyitrai and Geeves, 2004). The presence of the initial lag phase indicates that the formation of the open pocket conformation M(ADP) is not in rapid equilibrium (i.e. *k*_-1D_ >> *k*_+2D_).

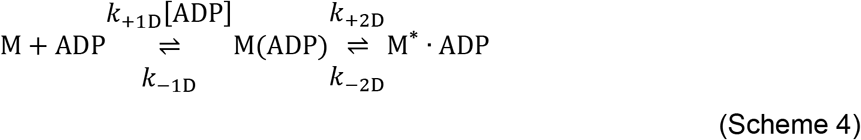

**FIGURE 6.**
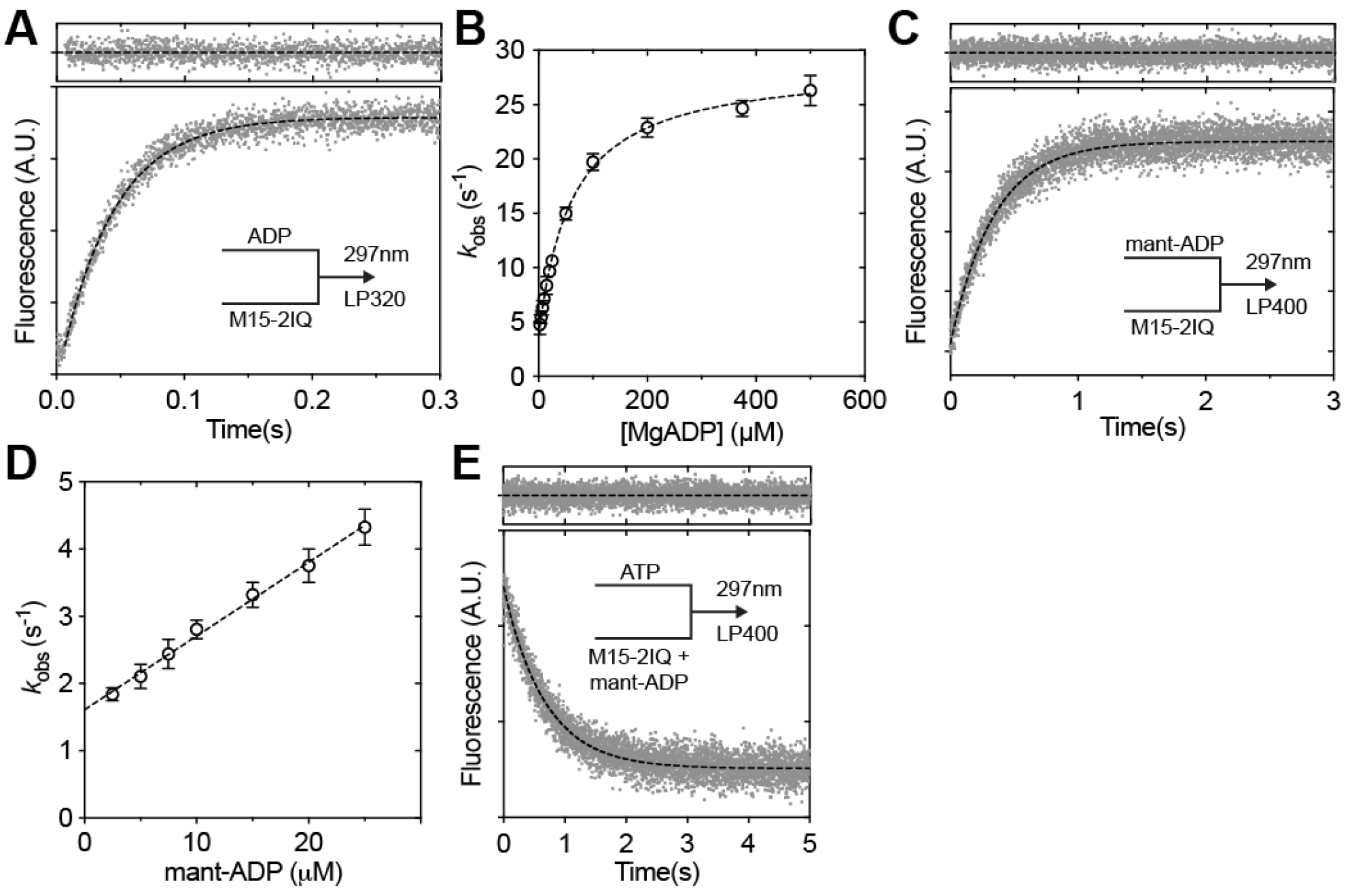
Kinetics of ADP binding to M15-2IQ. **A)** Enhancement of intrinsic protein fluorescence as 200 μM MgADP binds to 0.25 μM M15-2IQ in the stopped-flow. Following an initial lag phase (~ 10 ms), the recorded transient follows a single exponential time-course with *k*_obs_ = 23.8 ± 0.2 s^−1^ (dotted line, fit residuals above). **B)** Observed rate constants vary hyperbolically (dotted line) with respect to [MgADP], indicating that ADP binding is at least a two-step process. Referring to Scheme 4, the data points were fit to *k*_obs_ = *k*_-2D_ + (*K*_1D_*·k*_+2D_·[ADP]) / (1 + *K*_1D_·[ADP]), yielding *k*_+2_ = 25.2 ± 0.4 s^−1^, *k*_-2_ = 3.5 ± 0.3 s^−1^ and 1/*K*_1D_ = 60.6 ± 4.7 μM. **C)** Fluorescence enhancement following – 10 μM mant-ADP with 0.25 μM M15-2IQ in the stopped flow. The mant fluorophore was excited by FRET from vicinal tryptophan residues. The transient followed a mono-phasic exponential increase, *I(t) = −1.7e^−28.8t^ + C* (dotted line, fit residuals above). **D)** Observed rate constants varied linearly with [mant-ADP]. Linear regression to *k*_obs_ = *k*_+5_ + *k*_-5_ [ADP], yields *k*_-5_ = 0.11 ± 0.01 μM·s^−1^ and *k*_+5_ = 1.6 ± 0.07 s^−1^. **E)** Displacement reaction where 0.25 μM M15-2IQ was pre-equilibrated with 25 μM mant-ADP, before rapid mixing with 5 mM ATP. The transient followed a single exponential decay with *I(t) = 1.9e^−1.5t^ + C* (dotted line, fit residuals above). Reaction conditions for all experiments: 20 mM MOPS (pH 7.0), 100 mM KCl, 5 mM MgCl_2_, 0.1 mM EGTA at 20 °C. Data are representative of n = 3 independent determinations, except panel E (n = 2).

An analytical solution to this mechanism is a double exponential function with observed rate constants as defined in equations 1 and 2 (Hannemann et al., 2005). We fitted the slow observed rate constant (*k*_obs2_) from our data to equation 2, to yield estimates for the dissociation constant 1 / *K*+_1D_ = 60.6 ± 4.7 μM and isomerization rate constants *k*_+2D_ = 25.2 ± 0.4 s^−1^ and *k*_-2D_ = 3.5 ± 0.3 s^−1^ (Fig. 6B). The apparent association rate constant *K*_1D_ • *k*_+2D_ = 0.29 ± 0.04 μM^−1^s^−1^ was extracted from the gradient of the second-order response at low [ADP] (Fig. 6B). Although the initial binding of ADP (*K*_1D_) to M15-2IQ was low affinity (1 / *K*_1D_ ~ 60 μM), the overall apparent affinity is strongly influenced by *K*_2D_, the isomerization equilibrium constant (Nyitrai and Geeves, 2004). Equation 3 is an expression for the apparent dissociation equilibrium constant (*K*_D_) of a 2-step binding mechanism, rearranged from Nyitrai and Geeves (2004) to be consistent with Schemes 1 and 4. Equation 3 shows that if the isomerization *K*_2D_ << 1, then the apparent dissociation constant will equal the affinity of the open ADP-bound conformation, *K*_5_ = 1 / *K*1D. Conversely, if the isomerization *K*_2D_ >> 1, the apparent dissociation constant will be significantly tighter, with *K*_5_ = 1 / (*K*_1D_ • *K*_2D_). We calculated the isomerization equilibrium constant *K*_2D_ using the relation *K*_2D_ = *k*_+2D_ / *k*_-2D_ = 7.2 ± 0.6, and substituted into Equation 3 to yield the apparent dissociation constant *K*_5_ = 7.4 ± 0.8 μM.

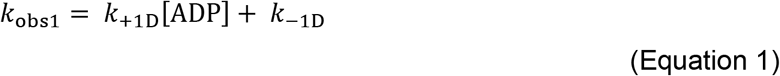

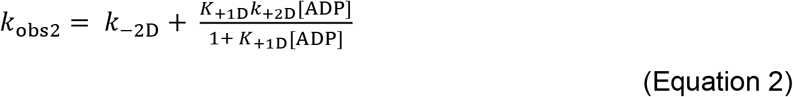

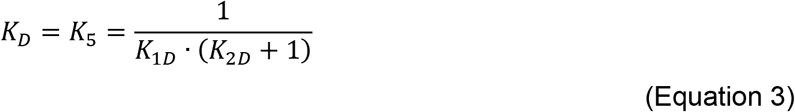

ADP binding to myosin was independently probed using the fluorescent analog, 2’-deoxy-mant-ADP (mant-ADP). M15-2IQ (0.25 μM) was rapidly mixed under pseudo-first order conditions in the stopped flow whilst titrating mant-ADP concentrations. The mant fluorophore was excited at 297 nm using FRET from vicinal tryptophan residues. Fluorescent transients followed a mono-phasic exponential time course with no lag phase (Fig. 6C). Observed rate constants (*k*_obs_) varied linearly with respect to [mant-ADP] and we calculated the apparent association rate constant *k*_-5_ = 0.11 ± 0.01 μM^−1^s^−1^ from the gradient of this second-order response, using the relation *k*_obs_ = *k*_-5_ • [ADP] + *k*_+5_ (Fig. 6D). The apparent dissociation constant *k*_+5_ = 1.6 ± 0.07 s^−1^ was measured from the y-axis intercept (Fig. 6D). A displacement reaction was used to independently confirm mant-ADP disassociation kinetics. M15-2IQ (0.25 μM) pre-equilibrated with 25 μM mant-ADP (> than *K*_d_ to ensure saturation of the active site) was rapidly mixed with 5 mM ATP in the stopped flow (Fig. 6E). The observed fluorescence decay followed a single exponential time course with *k*_obs_ = 1.50 ± 0.02 s^−1^. The dissociation equilibrium constant was calculated using the relationship, *K*_5_ = *k*_+5_ / *k*_-5_ = 13.6 ± 1.2 μM, in reasonable agreement with our estimate from intrinsic fluorescence.

We were unable to directly measure the kinetics of ADP binding to actomyosin using either intrinsic fluorescence or mant-labeled ADP analogs, as these transients had poor signal-to-noise. Instead, we indirectly probed the actin-attached ADP dissociation constant (*K’*_5_) by measuring ATP binding to actomyosin in the presence of ADP competing for the active site. We modelled this reaction according to Scheme 5, where ADP is in rapid equilibrium with ATP binding to actomyosin (De La Cruz and Ostap, 2009). M15-2IQ (0.125 μM) pre-equilibrated with actin (0.25 μM) and ADP, was rapidly mixed in the stopped-flow with 50 μM ATP, whilst measuring orthogonally scattered light to monitor dissolution of the actomyosin complex. The low concentration of ATP (50 μM, defined as ATP0) used in these titrations bound slowly to actomyosin (Fig. 3B) and allowed for ADP to compete in rapid equilibrium (i.e. *k’*_-5_ • [ADP] >> *K’*_1_ • *k*’_+2_ [ATP_0_]). Experimental transients were fit to a bi-phasic exponential decay in the absence of ADP (Fig. 7A), consistent with our earlier data that actomyosin occupies nucleotide-sensitive and insensitive states (Fig. 3B). Fit residuals showed evidence of a systematic deviation, but we could not consistently fit triple exponentials (Fig. 7A). Both fast and slow rate constants reduced hyperbolically as ADP was titrated from 0 – 500 μM (Fig. 7B). We focused on the observed fast phase (*k*_obs1_) and interpreted this as direct binding of ATP to the nucleotide-sensitive actomyosin fraction. Fast phase rate constants were fit to Equation 4, where ATP_0_ represents the initial [ATP] that is held constant throughout the titration (De La Cruz and Ostap, 2009). Fitting to the fast phase yielded the actin-attached ADP dissociation constant, *K’5* = 36.1 ± 5.7 μM. These data indicate a moderate thermodynamic coupling between the MYO15 nucleotide and actin binding sites (*K’*_5_ / *K*_5_ = 4.9), with the affinity of actomyosin for ADP (*K’*_5_) being 4.9-fold lower than the affinity of myosin for ADP (*K*_5_).

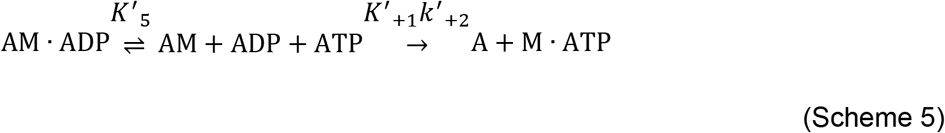

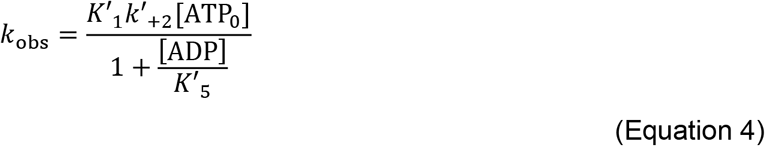

**FIGURE 7.**
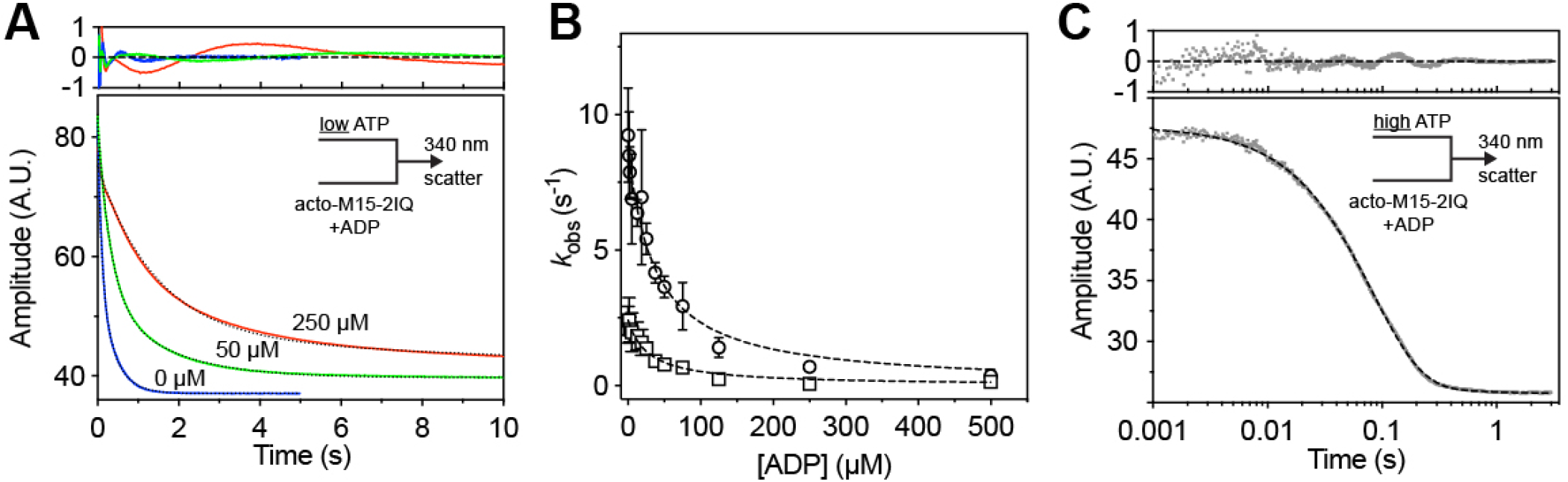
ADP binding to acto-M15-2IQ. **A)** The actin-attached dissociation constant (*K*’_5_) of ADP from M15-2IQ was measured using a displacement reaction. M15-2IQ (0.25 μM), actin (0.5 μM) and ADP (0 – 0.5 mM) were pre-equilibrated prior to rapid mixing with ATP (50 μM) in a stopped-flow spectrophotometer. ATP induced disassociation of the actomyosin complex was monitored using orthogonally scattered light at 340 nm. The ATP concentration was chosen to allow ATP and ADP to compete for binding to M15-2IQ. Example traces are shown for 0 μM (blue), 50 μM (green) and 250 μM (red) ADP, and are shown with bi-phasic exponential fits overlaid (dotted line, residuals above). For 250 μM ADP, the equation is defined by *I(t) = 28.3e^−40.2t^ + 6.6e^−0.1t^ + C*. **B)** Dependence of fast (circles) and slow (square) observed rate constants upon [ADP]. The fast phase was interpreted as direct binding of ATP to acto-M15-2IQ. Fitting of the fast-phase to an inverse-hyperbolae reveals half-maximal inhibition at 36.1 ± 5.7 μM ADP. **C)** ADP release from acto-M15-2IQ was measured using a displacement reaction. M15-2IQ (0.25 μM) was pre-equilibrated with actin (0.5 μM) and saturating ADP (0.5 mM), prior to rapid mixing with 5 mM MgATP in the stopped-flow. The high concentration of ATP prevents ADP rebinding. The light scattering transient is shown fit to a double exponential decay, *I(t) = 21.4e^−12.1t^ + 0.5e^−1.8t^ + C* (dotted line, residuals above). Reaction conditions for all experiments: 20 mM MOPS (pH 7.0), 100 mM KCl, 5 mM MgCl_2_, 0.1 mM EGTA at 20°C. Data are representative of n = 3 independent determinations, except panel C (n = 2).

In myosin motors that are specialized for processive motion along cellular actin filaments (i.e. MYO5A, MYO6), the rate limiting step of the ATPase cycle is actin-attached ADP release (*K’*_5_), ensuring that the motor has a high duty-ratio and remains attached to actin (De La Cruz et al., 1999, 2001). To determine if MYO15 has a similar characteristic, we measured actin-attached ADP release using a modification of the ADP competition experiment above. M15-2IQ (0.125 μM) was pre-equilibrated with actin (0.25 μM) and ADP (500 μM) to saturate the nucleotide binding site. The time course of light scattering was monitored following rapid mixing with 5 mM ATP in the stopped-flow. ATP binding to actomyosin was now rapid (~ 200 s^−1^) and irreversible under these conditions (Fig. 3B), and thus ATP binding was rate-limited by ADP release from the active site (Scheme 5). Experimentally observed light scattering transients were well fit to a double exponential transient with observed rate constants *k*_obs1_ = 12.5 ± 0.5 s^−1^ and *k*_obs2_ = 2.1 ± 0.5 s^−1^. The normalized amplitude of *k*_obs1_ was ~97% and we interpreted this to represent ADP release from actomyosin (*K’*_5_). Unlike MYO5A and MYO6 where the ATPase is rate-limited by actin-attached ADP release (De La Cruz et al., 1999, 2001), our data show that actin attached ADP release (~ 12 s^−1^) is not slow enough to solely rate-limit the MYO15 ATPase cycle (*k*_cat_ = 6 s-1).

## Discussion

### Client specificity of UNC45 chaperones for the MYO15 motor domain

This study expands upon our previous work that the striated-muscle UNC45B co-chaperone is necessary to promote folding of the MYO15 motor domain (Bird et al., 2014). In the previous study, we showed that expression of M15-2IQ in *Sf*9 cells resulted in aggregated, inactive protein unless UNC45B was co-expressed (Bird et al., 2014). In the present study we show the more ubiquitously expressed UNC45A has similar activity in *Sf*9 insect cells. UNC45A and UNC45B are co-chaperones that activate HSP90-dependent folding of the conventional muscle myosin motor domain (Barral et al., 2002; Liu et al., 2008), and also fold several unconventional myosin motor domains, including class XIV and XV (Bird et al., 2014; Bookwalter et al., 2014). Our experiments show that the MYO15 motor domain purified from *Sf*9 cells co-expressing either UNC45A or UNC45B, exhibit similar steady-state activities (*k*_cat_ and *K*_ATPase_), suggesting that the motor domains were functionally similar. These data do not support the hypothesis that different UNC45 chaperones post-translationally regulate motor domain activity, although we cannot rule out more subtle differences in ATPase activity that would be overlooked by steady-state assays. These experimental results used *Sf9* cells infected with a dual-promoter baculovirus expressing either mouse UNC45A or UNC45B, in conjunction with HSP90AA1. We further replicated these findings using Sf9 cells engineered to stably express only mouse UNC45A, or UNC45B, demonstrating that endogenous insect HSP90 orthologs were sufficient to function with UNC45. Our findings confirm a recent report where overexpression of the *C. elegans* UNC45 ortholog was sufficient to fold muscle myosin (MHC-B) in *Sf*9 cells without the overexpression of HSP90 (Hellerschmied et al., 2019). The *Sf*9-UNC45 cell lines developed here facilitate the expression of recombinant MYO15 and may prove helpful for other proteins and myosin motors that require UNC45 paralogs to fold and mature correctly.

Cochlear hair cells express both UNC45A and UNC45B (https://umgear.org) and we hypothesize that either one of these chaperones is required for the correct folding of MYO15 *in vivo*. Intriguingly, loss-of-function mutations in *UNC45A* are associated with a human form of syndromic deafness (Esteve et al., 2018), although it is unknown whether this is caused by defective hair cell stereocilia formation, similar to the phenotype caused by pathogenic *Myo15* variants (Probst et al., 1998; Anderson et al., 2000). Why might hair cells express both UNC45 paralogs? We suspect that the answer lies with UNC45 substrate specificity. HSP90-dependent folding of conventional smooth muscle myosin can be catalyzed *in vitro* by either UNC45 paralog, yet UNC45A displays increased folding efficiency towards this substrate (Liu et al., 2008). Whilst there appears to be some degree of functional overlap *in vitro*, developmental studies in zebrafish (*D. rerio*) show that *unc45a* and *unc45b* are not functionally redundant *in vivo* (Comyn and Pilgrim, 2012). We did not explore the relative efficiencies of UNC45A vs UNC45B in folding MYO15, however the expression of both paralogs points to UNC45 engaging with a broader spectrum of hair cell client proteins. A number of myosin genes are critical for hearing, including MYO3A / B (Walsh et al., 2011; Ebrahim et al., 2016; Lelli et al., 2016), MYO6 (Avraham et al., 1997), MYO7A (Liu et al., 1997) and MYO15 (Probst et al., 1998). MYO3A, MYO6 and MYO7A have all been successfully purified from *Sf9* cells without the use of UNC45 co-expression, suggesting that this chaperone is not a pre-requisite for folding (Wells et al., 1999; Dosé et al., 2008; Umeki et al., 2009; Yang et al., 2009). It is interesting to note that the UNC45 family chaperones also target the unfolded myosin motor domain *in vitro* (Barral et al., 2002; Srikakulam et al., 2008; Hellerschmied et al., 2019) and potentially do so in response to tissue trauma *in vivo* (Etard et al., 2008). An exciting hypothesis is that UNC45 paralogs may maintain hair cell proteostasis and refold myosin motors in stereocilia as they are denatured and damaged. Given that mammalian hair cells do not regenerate and must survive throughout the animal’s lifetime, we speculate this type of myosin quality control may be critical for long-lived sensory function.

### The ATPase mechanism of MYO15

Our experiments identify the key motor domain characteristics of MYO15 to be: 1) actin-attached ADP release as the slowest overall reaction step, 2) a moderate affinity for ADP with weak thermodynamic coupling between nucleotide and actin binding, 3) rapid ATP hydrolysis with an equilibrium constant (*K*_3_) close to unity, 4) weak binding of ATP, and 5) a moderate affinity for actin in the absence of nucleotide, compared to that of most myosins. Thermodynamic and kinetic rate constants from our steady-state and pre-steady state transient analyses are summarized in Tables II and III.

The basal ATPase rate of the MYO15 motor domain in the absence of actin (see Table II) was rate-limited by phosphate release (*k*_+4_), with the addition of actin stimulating the steady-state activity ~ 77-fold to a maximal turnover rate (*k*_cat_) of ~ 6 s^−1^ at 20°C, agreeing with previous estimates (Bird et al., 2014). We were unable to identify a single transition that solely rate-limited the catalytic cycle. Our transient kinetic analysis showed that ATP binding to actomyosin (*K’*_1_·*k’*_+2_) was > 125 s^−1^ at 2 mM MgATP, and thus too fast to be rate-limiting at steady state (Fig. 3B). Actin-detached hydrolysis (*k*_+3_ + *k*_-3_) was ~ 50 s^−1^ measured using quenched flow at 0.5 mM ATP, and was similarly too fast to be rate-limiting (Fig. 4A). We were unable to probe the maximum rate of actin-activated phosphate release (*k’*_+4_) due to the weak affinity of M·ADP·P_i_ for actin (*K*_9_) at the physiological salt concentrations (100 mM KCl) used in our study. Despite this, the second-order response was linear up to ~ 6 s^−1^ at 50 μM actin with no sign of deviation (Fig. 4C), indicating that phosphate release was not close to saturation, and unlikely to be rate-limiting either. ADP release from actomyosin was the slowest measured transition at ~ 12 s^−1^ and our study suggests is a major contributor towards rate-limiting the MYO15 ATPase cycle.

Both myosin and actomyosin had moderate affinities for ADP in the overall range of 7 – 36 μM, and thermodynamic coupling between nucleotide and actin binding was weak (*K*_AD_ / *K*_D_ = 2.7 – 4.9, *K*_DA_/*K*_A_ = 1.6, see Table III). This indicates that actin binding to MYO15 does not significantly lower its affinity for

ADP, and is a key enzymatic adaptation that allows for a load bearing, strongly bound actomyosin interaction to form under intracellular ADP concentrations (~ 50 μM) (Katz et al., 1989). This is in contrast to motors with high thermodynamic coupling ratios (*K*_AD_ / *K*_D_ > 30), such as fast skeletal myosin (Ritchie et al., 1993), where actin binding leads to a rapid release of ADP from the active site. Our data show that ADP binding and release from MYO15 involved at least a 2-step mechanism. We observed both a lag phase and saturation of observed rate constants as ADP bound to MYO15 (Fig. 6A,B). These data are consistent with a 2-step sequential binding mechanism and suggest that MYO15 undergoes an ADP-bound structural isomerization. Two-step ADP binding is proposed as a common mechanism for all hydrolysis-competent myosin motors, where ADP binds the nucleotide pocket in an ‘open’ configuration, before isomerization to an ADP-bound ‘closed’ configuration (Nyitrai and Geeves, 2004; Bloemink and Geeves, 2011). When strongly bound to actin, a critical parameter determining myosin function is the rate of isomerization, from the ADP-bound closed, to open isomer (also referred to as *K*_CO_). This isomerization can be opposed by mechanical load exerted on the actomyosin cross bridge, and modulate the rate of ADP release and thus the lifetime of the strongly-bound actomyosin state (Purcell et al., 2005; Veigel et al., 2005; Laakso et al., 2008; Oguchi et al., 2008).

We were unable to experimentally probe the kinetics of ADP-bound isomerization for actomyosin, as neither ADP nor mant-ADP generated robust fluorescence transients upon binding. Instead, we measured ADP release from actomyosin using a displacement reaction and observed a predominantly mono-phasic exponential decay of ~12 s^−1^ (Fig. 7C). A mono-phasic transient might be expected from a 2-step mechanism if the nucleotide binding pocket is biased toward the open ADP-bound configuration, since the closed ADP-bound species are unable to populate in the experimental pre-mixtures. If this were the case, our displacement reaction measured ADP release primarily from the open isomer, and it follows that any ADP-bound isomerization will be kinetically invisible in our experiment. ADP release from the closed isomer may only be kinetically accessible approaching from the hydrolysis and phosphate release states, similar to muscle myosin (Sleep and Hutton, 1980). In further support of an actin-attached ADP isomerization, we observed that ATP binding to nucleotide-free actomyosin was bi-phasic, with both phases saturating with increasing [ATP] (Fig. 3B). This behavior has been observed in class I myosin, and is consistent with the nucleotide-binding pocket of actomyosin isomerizing between open and closed states (Geeves et al., 2000; Lewis et al., 2006).

Similar to the other MyTH4-FERM myosins (MYO7A, MYO10), quenched flow experiments show that the MYO15 motor domain has a hydrolysis equilibrium constant (*K*_3_) near unity, indicating that M·ATP and M·ADP·P_i_ states are equally populated at steady-state (Kovács et al., 2005; Haithcock et al., 2011). This enzymatic adaptation may allow for a parallel flux through the actin-attached hydrolysis (i.e. M·ATP > AM·ATP· > AM·ADP·P_i_) pathway, in addition to actin-detached hydrolysis (i.e. M·ATP > M·ADP·P_i_ > AM·ADP·P_i_). The extent of this contribution depends upon the flux of actin-attached hydrolysis (*k’*_+3_ + *k’*_-3_) and the affinity of M·ATP for actin (*K*_8_, see Scheme 1). The latter is challenging to measure experimentally and has been previously estimated from global numerical simulations of the ATPase cycle (Kovács et al., 2005; Haithcock et al., 2011). In the case of MYO7A and MYO10, the effect of the actin-attached pathway is to divert enzymatic flux through a slower hydrolysis pathway and contribute to rate-limiting the ATPase cycle at higher actin concentrations (Kovács et al., 2005; Haithcock et al., 2011). It remains unclear from our study the extent of this contribution, if any, to the MYO15 ATPase cycle.

**Scheme 1.**
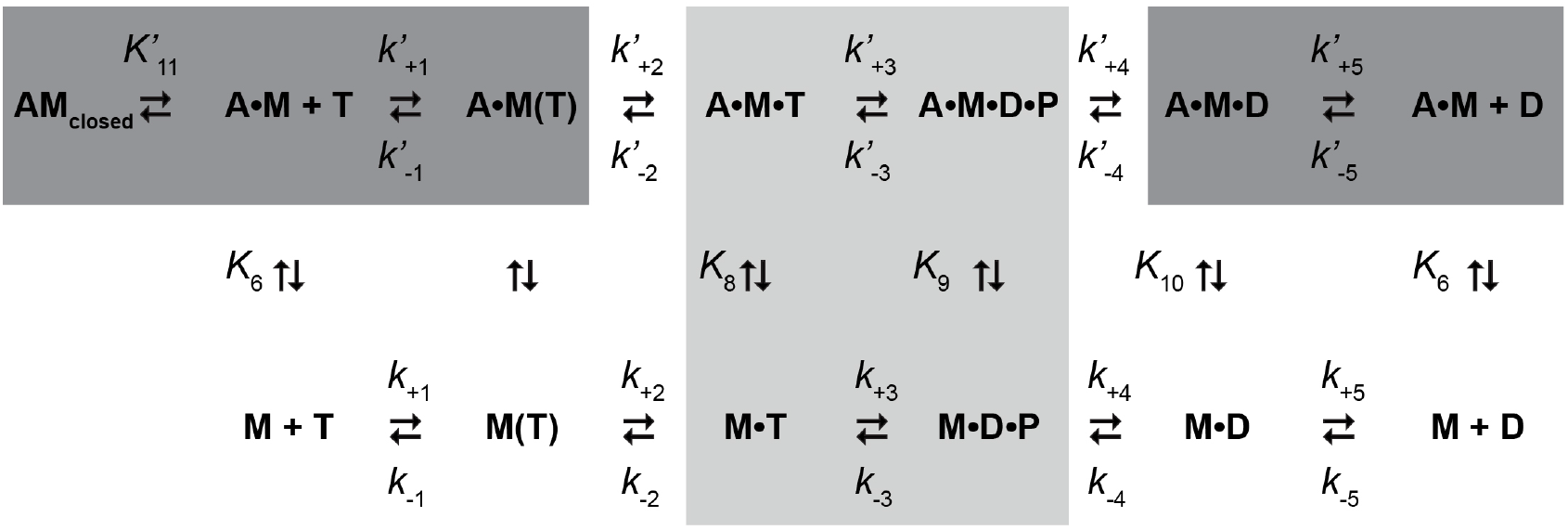
ATPase mechanism of the MYO15 motor domain. Species are abbreviated as follows: actin (A), myosin (M), ATP (T), ADP (D) and inorganic phosphate (P). Strongly actin-bound species (dark grey), weakly actin-bound species (light grey). Equilibrium constants are defined as *K*_i_ = *k*_+i_/ *k*_-i_, where the *k*_+i_ reaction proceeds up, or to the right. Actin-bound state transitions are denoted with an apostrophe (e.g. *k*’_+2_).

ATP binding to the MYO15 motor domain was notably weak with the rate constant not fully saturated at 5 mM ATP (Fig. 2B, 3B). This arises from the initial binding step having a high dissociation constant (1 / *K’*_1_ = 2.7 mM), and not from the subsequent isomerization step (*k’*+2) that was rapid (~ 300 s^−1^) and irreversible. As a consequence of the weak and slow binding of ATP, a proportion of the motor domain is expected to be in the nucleotide-free (rigor) state under physiological ATP concentrations. The weak binding of ATP suggests there may be steric hindrance either in the nucleotide binding pocket or gaining access to it. MYO6 exhibits even weaker ATP binding than MYO15 (1 / *K’*_1_ > 14 mM), and this has been attributed to a unique insert 1 in the nucleotide binding pocket that allows ADP to bind preferentially over ATP (De La Cruz et al., 2001; Pylypenko et al., 2011). MYO15 does not have an equivalent insert, however the motor domain structure has not been reported and it may reveal other steric effects in the binding pocket to explain this behavior. A possibility is that variations in loop 1 may contribute to the weak binding of ATP. MYO15 has an alternatively spliced cassette exon 8 that inserts 2 amino acids (“IK”) into loop one (Liang et al., 1999), and this may interfere with access to the pocket, or propagate changes to the nucleotide binding site via altered flexibility (Sweeney et al., 1998; Clark et al., 2005). M15-2IQ motor domain purified for this study contains the exon 8 encoded I and K residues, and it will be interesting to characterize the kinetics of the motor domain lacking these two residues.

The affinity of MYO15 for actin, either ADP bound (*K*_10_), or nucleotide free (*K*_6_) was relatively low, in the range of several hundred nM, compared with the high affinity of MYO5A / MYO6, for example (De La Cruz et al., 1999, 2001). The lower affinity results from a significant off-rate from actin (*k*_-6_, *k*_-10_) and is a characteristic shared with MYO10 which is related phylogenetically (Homma and Ikebe, 2005; Kovács et al., 2005; Odronitz and Kollmar, 2007). MYO10 exhibits only a small fractional quench of pyrene-labeled actin and is proposed to engage a subtly different actin binding interface from other myosins (Kovács et al., 2005). Cryo-EM reconstructions of the MYO10 motor domain at 9Å resolution have yet to reveal a structural basis for this (Ropars et al., 2016). The different actin binding topology may relate to MYO10 being kinetically optimized to move along bundles of actin filaments, in preference to individual filaments (Nagy et al., 2008; Ricca and Rock, 2010; Ropars et al., 2016). We expect MYO15 to also be specialized for trafficking along bundles of actin filaments, given that stereocilia actin filaments are extensively cross-linked with FSCN1/2, PLS1, ESPN and TRIOBP (Bretscher, 1981; Zheng et al., 2000; Kitajiri et al., 2010). If this is true, we may expect some kinetic parameters for MYO15 to be altered when measured on bundled actin versus single actin filaments, similar to MYO10 (Ropars et al., 2016).

### Duty ratio of MYO15

The duty ratio of a myosin molecular motor is defined as the fraction of its mechano-chemical cycle spent attached to the actin filament (Howard, 1997). Under saturating ATP, the AM (apo) state is not significantly occupied at steady-state, and thus the duration spent attached to actin is dominated by the strongly bound AM•ADP state. The duty ratio can be expressed as follows (Equation 5).

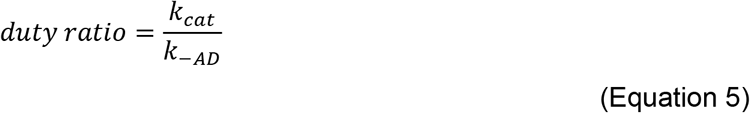

Using our measured values for *k*_cat_ and *k*_-AD_ (*k*’_+5_), we estimate that MYO15 has a duty ratio = 0.48 indicating that the motor domain spends ~ 48% of its cycle strongly bound to an actin filament. This calculation assumes that entry to the AM•ADP state is not rate-limited by either ATP binding (*k’*_+2_), ATP hydrolysis (*K*_3_), or actin-attached phosphate release (*K*_9_•*k’*_+4_), and omits the dependence of duty ratio upon [actin]. The duty ratio calculated above would be achieved at a hypothetical infinite [actin], and we speculate this may be biologically relevant considering that MYO15 operates within the densely packed actin filaments of the stereocilia actin core. The duty ratio calculated here differs from our previous estimate of > 0.9 that was inferred from the dwell time of actin-attached binding events in an optical trap (Bird et al., 2014). Uncertainties in the 2-step ADP release mechanism from actomyosin may account for this difference; an ADP isomerization step (*K*_CO_) slower than 12 s^−1^ would increase the steady-state population of myosin in strongly actin-bound states, and thus the duty ratio. Furthermore, if ADP release from actomyosin is force sensitive, such as shown for MYO1B (Laakso et al., 2008), then the low pN forces experienced in the optical trap may retard ADP release and further increase the duty ratio. Future experiments using force-feedback optical trapping are needed to test if ADP release from MYO15 is load dependent, and if so whether a dimerized molecule could gate ADP-release to further increase overall processivity, similar to MYO5A (Purcell et al., 2005; Veigel et al., 2005). For these reasons, we consider the duty-ratio estimate of 0.48 in our present study to be a lower bound.

### Implications for MYO15 function within hair cells

The cellular functions of myosin motors have been classified into four broad overlapping categories, based upon the kinetic signature of their motor domains (De La Cruz and Ostap, 2004; Nyitrai and Geeves, 2004; Bloemink and Geeves, 2011). This scheme utilizes four motor domain parameters: 1) duty ratio, 2) thermodynamic coupling of actin and nucleotide binding (*K*_AD_ / *K*_D_), 3) load dependence of ADP release, and 4) *K*_C/O_, the equilibrium constant of actin-attached ADP isomerization (from closed to open nucleotide pocket). The kinetic adaptations revealed in our present study show that MYO15 has an intermediate duty ratio (~ 0.5) in addition to low thermodynamic coupling between nucleotide and actin binding sites. Combined, these characteristics enable the MYO15 motor domain to spend at least 50% of its mechanochemical cycle attached to actin, and to simultaneously bind actin and ADP allowing for a force-bearing cross bridge to form. These characteristics are consistent with a monomeric MYO15 molecule behaving as a strain sensor, where force applied to the motor domain can modulate ADP release and its actin-attached lifetime. If MYO15 molecules were to oligomerize into an ensemble, these same characteristics could also give rise to kinetic gating between motor domains to allow for longer-distance processive movement along an actin filament.

There is significant evidence that MYO15 molecules can move processively *in vivo*. MYO15 isoforms accumulate at the distal tip of hair cell stereocilia (Belyantseva et al., 2003; Fang et al., 2015) and time-lapse studies show a continual flux of motors towards the stereocilia tip (Hwang et al., 2015). Furthermore, EGFP-tagged MYO15-2 (Fig. 1A, isoform 2) traffics towards the distal tips of actin-based filopodia in COS-7 cells (Belyantseva et al., 2003), in a striking parallel with MYO10 (Berg and Cheney, 2002). MYO15-2 does not appear to move anterogradely in large puncta along filopodia (Belyantseva et al., 2005), suggesting that they instead move in small packets consisting of a few molecules, similar to MYO10 (Kerber et al., 2009). Consistent with this, small packets of MYO15 are observed in-transit along the stereocilia shaft in paraformaldehyde-fixed hair cells (Rzadzinska et al., 2004). With an intermediate duty ratio (~ 0.5), our kinetic study predicts that MYO15 needs to oligomerize in order to move processively within stereocilia. How this might be achieved is unclear. Whilst MYO15 lacks any coiled-coil motifs, it may use a cargo-mediated mechanism similar to MYO6 and MYO7A, where accessory proteins activate motility by driving oligomerization of the myosin heavy chain (Spudich et al., 2007; Phichith et al., 2009; Sakai et al., 2011). MYO15 binds at least four proteins in hair cells, including WHRN, EPS8, GNAI3 and GPSM2. These proteins are trafficked as a complex by MYO15 and regulate stereocilia elongation (Belyantseva et al., 2005; Delprat et al., 2005; Manor et al., 2011; Mauriac et al., 2017; Tadenev et al., 2019). We speculate that an additional function of these proteins, or a hitherto unidentified partner, is to oligomerize MYO15 and to activate processive motility towards the stereocilia tip.

In addition to trafficking within stereocilia, MYO15 concentrates at the stereocilia tip where active mechano-electric transduction (MET) channels are located (Beurg et al., 2009). Recent studies have shown that the MYO15:WHRN complex binds via CIB2 (Riazuddin et al., 2012; Giese et al., 2017) to TMC1 and TMC2; pore-forming subunits of the MET channel (Kawashima et al., 2011; Pan et al., 2018).

Our finding that the MYO15 motor domain is kinetically tuned as a strain sensor suggests that MYO15 could also act as a force-sensitive element bridging the membrane and actin cytoskeleton at the stereocilia tip. Whilst the physiological function of this remains speculative, MYO15 is ideally placed to respond to tension exerted during auditory mechano-transduction. Understanding how MYO15 oligomerizes *in vivo* is a key question for future experiments, with the monomer to oligomer transition potentially controlling the switch between force-sensing and processive activity.

Our kinetic study represents an important step towards understanding how pathogenic mutations in *MYO15A* cause human deafness DFNB3. More than 300 mutant alleles of *MYO15A* have been identified that cause DFNB3, and a significant percentage of these are missense mutations substituting residues in the motor domain (Rehman et al., 2016). The effects of these mutations are unknown. Our detailed study provides a thermodynamic and kinetic “fingerprint” of the wild-type motor domain, and will allow the precise effects of deafness causing mutations to be identified. Our results will also help reveal the activity of full-length MYO15 isoforms that have distinct cellular functions in the assembly and maintenance of the stereocilia actin core (Fang et al., 2015; Rehman et al., 2016). MYO15 isoforms are distinguished by a N-terminal domain that precedes the motor domain (Fig. 1A). The N-terminal domain of MYO15-1 is notable for both its size (~1200 aa) and unusually high proline content (17%). In class I myosins, the artificial switching of N-terminal domains between the MYO1B and MYO1C motor domain enables strain-sensitive binding to actin filaments (Laakso et al., 2008; Shuman et al., 2014; Greenberg et al., 2015; Mentes et al., 2018). We hypothesize that the N-terminal domain may similarly regulate the motor activity of MYO15-1, and endow this molecule with unique properties. By comparison to the motor domain activity reported in our present study, we hope to understand how the enzymatic activities of full-length MYO15 isoforms differ, and ultimately how they contribute to the stereocilia development and maintenance processes essential for lifelong hearing.

### Experimental procedures

#### General reagents

Reagents were of the highest grade available from Millipore Sigma (St Louis, MO), unless otherwise stated. ATP and ADP nucleotides were prepared as equimolar stocks with magnesium acetate at pH 7.0 and quantified by absorbance at 259 nm (ε = 15,400 M^−1^·cm^−1^). 2’deoxy-mant labeled ATP and ADP stocks (BIOLOG, Germany) were quantified by absorbance at 255 nm (ε = 23,300 M^−1^·cm^−1^). Coumarin labeled phosphate-binding protein (MDCC-PbP)(Brune et al., 1994) was the generous gift of Dr. Howard White (University of Virginia Medical School). ATP [gamma-32P] was from Perkin Elmer (Waltham, MA).

#### Actin purification and labeling

Rabbit skeletal actin was extracted from muscle acetone powder (Pel-Freez, AZ) and labeled on Cys-374 using *N*-(1-pyrene)-iodoacetamide (Thermo Fisher Scientific) as needed (Spudich and Watt, 1971; Criddle et al., 1985). Actin for steady-state ATPase measurements was purified through two rounds of polymerization / depolymerization with ultracentrifugation. The concentration of polymerized actin was determined by measuring the absorbance at 290 nm (ε = 26,600 M^−1^ cm^−1^) and dialyzed extensively against 4 mM MOPS (pH 7.0), 1 mM MgCl_2_, 0.1 mM EGTA, 1 mM DTT, 1 mM NaN3 prior to use. All actins used for stopped-flow spectrophotometry (unlabeled and pyrene-conjugated) were purified by size exclusion chromatography (16/60 Sephacryl S-300 HR, Purifier 10, GE Healthcare) with isocratic elution in G-buffer (2 mM Tris HCl (pH 8.0), 0.2 mM ATP, 0.1 mM CaCl2, 1 mM DTT), prior to polymerization and extensive dialysis against 20 mM MOPS (pH 7.0), 100 mM KCl, 5 mM MgCl_2_, 0.1 mM EGTA, 1 mM DTT to remove contaminating nucleotides. Actin concentrations were determined at 290 nm (ε = 26,600 M^−1^ cm^−1^) and additionally at 344 nm (ε = 22,000 M^−1^ cm^−1^) for pyrene-labeled stocks. A correction factor was applied for pyrene actin, A_corr_ = A_290_ – 0.127 * A_344_. Pyrene labeling of actin was typically 80 - 90% (mol : mol). All actin filaments used for stopped-flow experiments were stabilized with a molar equivalent of phalloidin (Sigma).

#### *Sf*9-UNC45 stable cell generation and western blotting

*Cloning of pIB-Unc45a and pIB-Unc45b:* Transient expression of untagged UNC45A or UNC45B in *Sf9* cells was driven using the opIE2 promoter in *pIB* (Thermo Fisher Scientific). Expression of *pIB* confers resistance to the antibiotic blasticidin allowing for positive selection in *Sf9* insect cells. The *pIB* empty vector was linearized with EcoRI and XhoI prior to agarose electrophoresis and gel purification (NucleoSpin, Takara Bio). The open reading frames encoding mouse UNC45A and mouse UNC45B were PCR amplified from *pFastbac Dual Unc45(a /b) /Hsp90aa1* (see below) using PrimeStar HS (Takara Bio, CA) and primers as shown (Table I). A stop codon was introduced into the reverse primers to prevent expression of the V5-His6 tag within the *pIB* vector backbone. Amplicons were ligated into linearized *pIB* using InFusion HD EcoDry (Takara Bio) and transformed into *E.coli* bacteria (Stellar, Takara Bio) following the manufacturer’s protocol. Recombinant clones were screened by restriction digest with EcoRI and XhoI, and correctly structured plasmids were Sanger sequenced to confirm the open reading frame.

**Table I.**
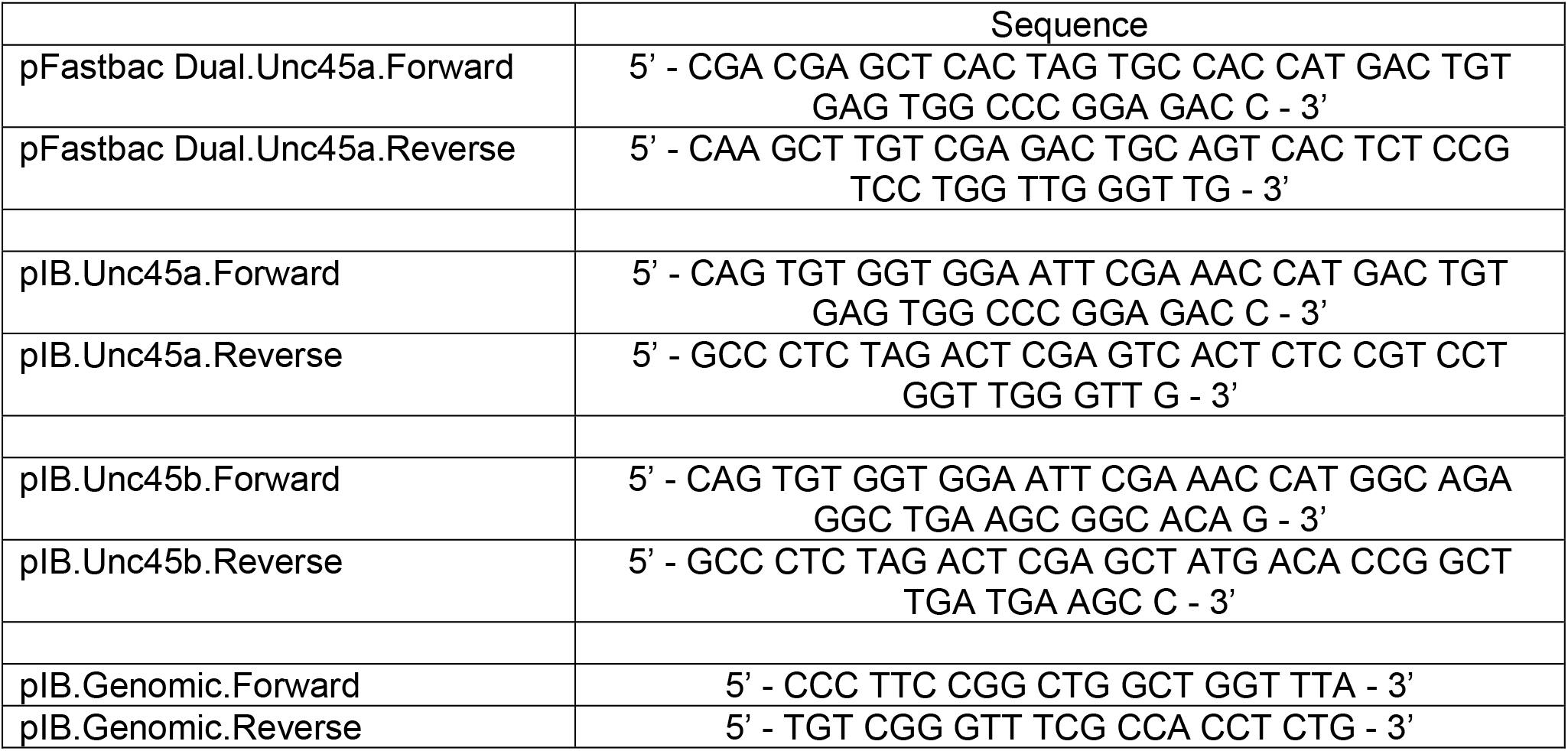
(DNA oligonucleotides used in this study)

**Table II.** (Summary of steady-state ATPase measurements)

| | $k_{\text{cat}}$ ($\text{s}^{-1}$) | $k_{\text{ATPase}}$ ($\mu\text{M}$) | Basal ATPase ($\text{s}^{-1}$) |
| --- | --- | --- | --- |
| M15-2IQ (baculovirus UNC45A) | $5.6 \pm 0.1$ | $6.1 \pm 0.5$ | $0.05 \pm 0.01$ |
| M15-2IQ (baculovirus UNC45B) | $5.9 \pm 0.4$ | $10.4 \pm 2.2$ | $0.06 \pm 0.01$ |
| M15-2IQ-EGFP (Sf9 - UNC45A) | $6.0 \pm 0.5$ | $5.2 \pm 1.5$ | $0.1 \pm 0.01$ |
| M15-2IQ-EGFP (Sf9 - UNC45B) | $5.8 \pm 0.3$ | $7.0 \pm 1.3$ | $0.09 \pm 0.01$ |

Experimental conditions: 10 mM MOPS pH 7.0, 10 mM KCl, 5 mM $\text{MgCl}_2$ , 0.1 mM EGTA, 2 mM MgATP at $20 \pm 0.1$ °C. Data are mean ± SEM (for $k_{\text{cat}}$ and $k_{\text{ATPase}}$ ) and mean ± SD (for basal ATPase).

**Table III.** (Summary of key ATPase kinetic parameters)

| **ATP binding** |  |
| --- | --- |
| $K_1 \cdot k_{+2} (\mu\text{M}^{-1} \cdot \text{s}^{-1})$ | $0.17 \pm 0.01$ intrinsic<br>$0.15 \pm 0.002$ mant ATP |
| $1 / K_1 (\mu\text{M})$ | $1898 \pm 145$ intrinsic |
| $k_{+2} (\text{s}^{-1})$ | $322 \pm 10$ intrinsic |
| $K'_1 \cdot k'_{+2} (\mu\text{M}^{-1} \cdot \text{s}^{-1})$ | $0.12 \pm 0.02$ light scatter<br>$0.12 \pm 0.01$ mant ATP |
| $1 / K'_1 (\mu\text{M})$ | $2749 \pm 268$ light scatter |
| $k'_{+2} (\text{s}^{-1})$ | $307 \pm 15$ light scatter |
| **ADP binding** |  |
| $k_{+5} (\text{s}^{-1})^{\text{app}}$ | $2.1 \pm 0.4 (= K_D \cdot k_{-5})$ intrinsic<br>$1.5 \pm 0.02 \text{ s}^{-1}$ mant ADP |
| $k_{-5} (\mu\text{M}^{-1} \cdot \text{s}^{-1})^{\text{app}}$ | $0.29 \pm 0.04$ intrinsic<br>$0.11 \pm 0.01$ mant ADP |
| $K_5 = K_D (\mu\text{M})^{\text{app}}$ | $7.4 \pm 0.8$ intrinsic<br>$13.6 \pm 1.2$ mant ADP |
| $k'_{+5} (\text{s}^{-1})^{\text{app}}$ | $12.5 \pm 0.5$ light scatter |
| $k'_{-5} (\mu\text{M}^{-1} \cdot \text{s}^{-1})^{\text{app}}$ | $0.35 \pm 0.06 (= k'_{+5} / K'_5)$ light scatter |
| $K'_5 = K_{\text{AD}} (\mu\text{M})^{\text{app}}$ | $36.1 \pm 5.7 \mu\text{M}$ light scatter |
| $K_{\text{AD}} / K_D$ | $4.9 (= K'_5 / K_5^{\text{intrinsic}})$<br>$2.7 (= K'_5 / K_5^{\text{mant ADP}})$ |
| **Hydrolysis and phosphate release** |  |
| $k_{+3} + k_{-3} (\text{s}^{-1})$ | $322 \pm 10$ intrinsic<br>$\geq 50.2 \pm 5.7$ quenched flow |
| $K_3$ | $1.34 \pm 0.07$ quenched flow |
| $k_{+4} (\text{s}^{-1})$ | $0.08 \pm 0.001$ MDCC-PbP |
| $K_9 \cdot k'_{+4} (\mu\text{M}^{-1} \cdot \text{s}^{-1})$ | $0.13 \pm 0.004$ MDCC-PbP |
| **Actin binding** |  |
| $k_{+6} (\mu\text{M}^{-1} \cdot \text{s}^{-1})$ | $3.18 \pm 0.15$ pyrene |
| $k_{-6} (\text{s}^{-1})$ | $0.65 \pm 0.3$ pyrene <sup>#</sup> |
| $1/K_6 = K_A (\text{nM})$ | $200 \pm 100$ pyrene |
| $k_{+10} (\mu\text{M}^{-1} \cdot \text{s}^{-1})$ | $1.75 \pm 0.09$ pyrene |
| $k_{-10} (\text{s}^{-1})$ | $0.55 \pm 0.17$ pyrene |
| $1/K_{10} = K_{\text{DA}} (\text{nM})$ | $310 \pm 97$ pyrene |
| $K_{\text{DA}} / K_A$ | 1.6 |
| **Other** |  |
| $K'_{11}$ | $3.8 \pm 1.0$ light scatter |
| $k'_{+11} (\text{s}^{-1})$ | $30.9 \pm 1.9 \text{ s}^{-1}$ light scatter |
| $k'_{-11} (\text{s}^{-1})$ | $8.1 \pm 2.2 (= k'_{+11} / K'_{11})$ |

Assay conditions: 20 mM MOPS pH 7.0, 100 mM KCl, 5 mM MgCl<sub>2</sub>, 0.1 mM EGTA at 20 ± 0.1 °C.
<sup>app</sup> apparent values; <sup>#</sup> inferred from y-axis intercept.

*Sf9* cells were maintained at 27°C in HyClone SFX (GE Healthcare) and transfected in suspension culture using endotoxin-free *pIB-Unc45a*, or *pIB-Unc45b* plasmid DNA (NucleoBond Maxi EF, Takara Bio). Plasmid DNA was complexed with polyethylenimine (PEI, #24765-1, Polysciences Inc., PA) at a 12:1 (w/w) PEI:DNA ratio and added to *Sf9* cells in suspension culture. After 96 hours in culture, cells were sparsely seeded into 10 cm culture dishes and media supplemented with 15 μg/mL blasticidin S (Thermo Fisher Scientific) to drive positive selection of *Sf*9 cells with stable integration of the *pIB-Unc45a*, or *pIB-Unc45b* plasmid. After approximately 2 weeks in culture, adherent colonies were isolated using glass cloning cylinders (Sigma), transferred to a 96-well plate, and sequentially scaled up to a 6-well plate. The identity of each clone was analyzed by genomic PCR and western blotting. The *pIB* gene expression cassette containing *Unc45a* or *Unc45b* was PCR amplified from Sf9 genomic DNA using ExTaq (Takara Bio) and primers as shown (Table I). Amplicons of the expected size (~7.5 kb) were gel purified and Sanger sequenced to confirm integrity of the UNC45 transgene. Soluble protein was extracted from *Sf*9-UNC45A and *Sf*9-UNC45B cells using lysis buffer (see below: Expression and purification of M15-2IQ), analyzed by SDS-PAGE (4-20% TGX, BIO-RAD) and blotted to PVDF membrane using semi-dry transfer (Turbo-Blot, BIO-RAD). Blots were probed with primary antibodies, mouse IgG anti-UNC45A (SAB1400633, Millipore Sigma) or mouse IgG anti-UNC45B (ab77062, Abcam, MA), followed by secondary detection with horseradish peroxidase conjugated goat anti-mouse IgG (AP308P, Millipore Sigma). Chemi-luminescence was captured using a charge-coupled camera (ChemiDoc MP, BIO-RAD).

#### Baculoviral transfer vector cloning and baculovirus generation

A dual promoter baculoviral vector (*pFastbac Dual*, Life Technologies, CA) was used to engineer baculovirus expressing either mouse UNC45A (GenBank: AAH04717.1) or UNC45B (GenBank: AAH84585.1), in addition to mouse HSP90AA1 (GenBank: AAH46614.1) from the polyhedrin (PH) and p10 promoters, respectively. UNC45A, UNC45B and HSP90AA1 expressed proteins were not epitope tagged. The generation of *pFastbac Dual Unc45b* / *Hsp90aa1* was previously described (Bird et al., 2014). To generate *pFastbac Dual Unc45a* / *Hsp90aa1*, the open reading frame (ORF) of mouse *Unc45a* was PCR amplified from a P8.5 mouse embryo cDNA (Takara Bio, CA) using PrimeStar HS (Takara Bio) and primers as shown (Table I). Amplicons were ligated into SpeI / PstI linearized *pFastbac Dual Hsp90aa1* (Bird et al., 2014) using InFusion HD EcoDry (Takara Bio). Correctly recombined clones were isolated and confirmed by Sanger sequencing of both *Unc45a* and *Hsp90aa1* ORFs. *pFastbac1 EGFP-M15-2IQ* and *pFastbac M15-2IQ-EGFP* encoding the truncated mouse MYO15 motor domain (NP_874357.2, aa 1 - 743) with two LCBDs (IQ domains) and a C-terminal FLAG (DYKDDDK) epitope, have been previously described (Bird et al., 2014). The expressed proteins were (His)_6_-tev-EGFP-M15-2IQ-FLAG (119 kDa) and M15-2IQ-EGFP-FLAG (114 kDa), respectively.

Baculoviral transfer vectors were transformed into *E.coli* DH10-Bac cells (Life Technologies, CA), and recombinant bacmid DNA purified according to the manufacturer’s protocol. Recombinant bacmid DNA was transfected into *Sf9* cells using PEI, as described above, to generate first passage (P1) baculoviral stocks. A dual promoter baculovirus encoding bovine smooth muscle essential (MLC17B, MYL6) and chicken regulatory (MLC20, MYL12B) light chains was previously described (Pato et al., 1996). First passage baculoviral stocks were amplified in *Sf9* cells, using a low multiplicity of infection (MOI = 0.1), to generate second passage (P2) virus for expression. All baculoviral stocks were titered using an end-point dilution assay in combination with the Sf9-ET (Easy Titer) reporter cell line (Hopkins and Esposito, 2009).

#### Expression and purification of M15-2IQ

*Sf9* cells were maintained in either HyClone SFX (GE Healthcare) or ESF-921 (Expression Systems, CA) at 27°C in suspension culture. For protein expression, cells were seeded at 2 x 10^6^ cells·mL^−1^ and infected simultaneously (MOI = 5) with three baculoviruses encoding M15-2IQ, UNC45(A / B) / HSP90AA1 and light-chains (ELC / RLC). Infected cells were harvested after 48 hours and flash frozen in liquid nitrogen.

Purification of M15-2IQ was performed as previously described (Bird et al., 2014). Briefly, *Sf9* cell pellets were thawed in 10 mM MOPS (pH 7.2), 0.5 M NaCl, 1 mM EGTA, 10 mM MgCl_2_, 2 mM ATP, 0.1 mM DTT, 0.2 mM PMSF, 1 mM NaN3, 2 μg·mL^−1^ leupeptin and a protease inhibitor cocktail (Halt EDTA-free; Thermo Scientific) and lysed using a Dounce homogenizer chilled on ice. Sf9 cells lysates were sedimented at 48 kGmax × 30 min and the supernatant incubated for 3 hours on ice with FLAG-M2 affinity resin (Sigma-Aldrich). FLAG-M2 affinity resin was packed into a gravity-flow column and washed with high-salt buffer, 10 mM MOPS (pH 7.2), 0.5 M NaCl, 1 mM EGTA, 5 mM MgCl_2_, 1 mM ATP, 1 mM NaN3, 0.1 mM DTT, 0.1 mM PMSF, 1 μg·mL^−1^ leupeptin, followed by a low salt buffer, 10 mM MOPS (pH 7.0), 0.1 M NaCl, 1 mM EGTA, 1 mM NaN3, 0.1 mM DTT, 0.1 mM PMSF, 1 μg·mL^−1^ leupeptin. M15-2IQ protein was eluted from the FLAG affinity matrix using low-salt buffer supplemented with 0.2 mg·mL^−1^ 3X FLAG peptide (American Peptide, CA). Eluted M15-2IQ was bound to a strong anion exchanger (5/50 MonoQ GL; GE Healthcare) using a Purifier 10 (GE Healthcare) chromatography system at 4 °C. Bound protein was washed using 5 column volumes (CV) of 10 mM MOPS (pH 7.0), 0.1 M NaCl, 1 mM EGTA, 0.1 mM PMSF, 1 mM DTT and eluted with a 160 CV gradient to 1 M NaCl. M15-2IQ fractions eluting at ~31 mS·cm^−1^ were concentrated (10,000 MWCO; Amicon, Millipore) and further purified by size exclusion chromatography. M15-2IQ was loaded onto a HiLoad 16/60 column packed with Superdex 200 (GE Healthcare), and purified using isocratic elution with 10 mM MOPS (pH 7.0), 250 mM KCl, 0.1 mM EGTA, 1 mM NaN3, 0.1 mM PMSF, 1 mM DTT, 1 μg·mL^−1^ leupeptin, 10% (vol / vol) glycerol. M15-2IQ bound to ELC and RLC eluted as a single peak. The concentration of the final protein complex was determined at 280 nm using the following extinction coefficients, and assuming a 1 : 1 : 1 stoichiometry (EGFP-M15-2IQ : ELC : RLC, ε = 93,980 M^−1^·cm^−1^; M15-2IQ-EGFP : ELC : RLC, ε = 88,020 M^−1^·cm^−1^).

#### Transient and steady-state kinetic measurements

Steady-state ATPase activity was measured using a NADH coupled enzyme assay in the following reaction buffer: 10 mM MOPS (pH 7.0), 5 mM MgCl_2_, 0.1 mM EGTA, 10 mM KCl, 2 mM MgATP, 40 U·mL^−1^ lactate dehydrogenase, 200 U·mL^−1^ pyruvate kinase, 1 mM phosphoenolpyruvate, 200 μM NADH. The time course of NADH absorbance at A340 was measured using a UV-1800 dual-beam spectrophotometer (Shimadzu, MD) with a multi-cell cuvette at a constant 20 ± 0.1 °C. The rate of ATP consumption (i.e. ADP production) was calculated from the change in NADH using ε = 6,220 M^−1^·cm^−1^. The final concentration of myosin in each reaction was 30 nM.

Transient kinetic analysis was performed using a HiTech SF61-DX2 dual-mixing stopped-flow spectrophotometer (TgK Scientific, UK) or RQF-3 quenched-flow (Kintek, PA). Stopped-flow excitation wavelength and emission filters were as follows: intrinsic protein fluorescence (297 nm, WG320), pyrene-actin (365 nm, GG400), MDCC-PbP (430 nm, LP450) and mant-nucleotides (297 nm, GG400). Orthogonal light scattering was measured at 340 nm without emission filtering. Concentrated stocks of actin and myosin were pre-treated using 0.02 U/mL apyrase for 30 minutes at room temperature to remove trace ATP and ADP contamination. After dilution to the final experimental dilution, the concentration of apyrase was < 0.001 U·mL^−1^. For phosphate release experiments, the stopped flow apparatus and all solutions were incubated with 7-methylguanosine (0.5 mM) and nucleoside phosphorylase (0.02 U·mL^−1^) to scavenge contaminating Pi. The fluorescent phosphate-binding probe, MDCC-PbP, was included in all reactants at a final concentration of 5 μM. The volume ratio for single mixing stopped-flow reactions was 1:1. For double-mixing phosphate release measurements, ATP and M15-2IQ were mixed at a 1:1 ratio, aged in a delay loop, prior to mixing 1:1 with actin. The final ratio of ATP, M15-2IQ and actin in the observation cell was 1:1:2 (vol / vol), respectively. Quenched flow analysis was performed as previously described (Kovács et al., 2005). In general, all reactions were mixed under pseudo-first order conditions, with one reactant in > 10-fold excess, unless otherwise stated. Rate and equilibria constants were extracted from experimental data using established analytical approaches (Johnson, 1992; De La Cruz et al., 1999; De La Cruz and Ostap, 2009). All reactions were performed in 20 mM MOPS (pH 7.0), 100 mM KCl, 5 mM MgCl_2_, 0.1 mM EGTA at 20 ± 0.1 °C.

### Data analysis

Non-linear least-squares regression was performed using Kinetic Studio (TgK Scientific, UK) and Prism 8 (GraphPad). Uncertainties arising from non-linear regression are reported as standard error, and for all other measurements as standard deviation. Uncertainties were propagated for parameters that required multiplication or division of primary experimental values. Experimental data are from three independent experiments, using at least two independent myosin preparations, unless otherwise stated.

## Author contributions

Conceptualization: TBF, JRS, JEB

Formal analysis: FJ, YT, AS, SMH, JEB

Funding acquisition: JRS, TBF, JEB

Investigation: FJ, YT, AS, SMH, JEB

Project administration: JEB

Resources: JEB

Writing original draft: JEB

Writing – review and editing: FJ, YT, AS, SMH, TBF, JRS, JEB

## Data availability statement

All data referenced are contained within the article. The *Sf*9-UNC45A and *Sf*9-UNC45B cell lines are available from JEB.

## Acknowledgements

We are grateful to Howard White for the generous gift of purified MDCC-PbP, and thank Earl Homsher and Lee Sweeney for insightful discussions and critical feedback. This work was supported (in part) by the NIH Intramural Research Program of the NHLBI to JRS (HL006049), NIDCD to TBF (DC000048) and University of Florida startup funds to JEB.

